# Identification of the unwinding region in the *Clostridioides difficile* chromosomal origin of replication

**DOI:** 10.1101/2020.07.08.193532

**Authors:** Ana M. Oliveira Paiva, Erika van Eijk, Annemieke H. Friggen, Christoph Weigel, Wiep Klaas Smits

**Affiliations:** Department of Medical Microbiology, Section Experimental Bacteriology, Leiden University Medical Center, Leiden, The Netherlands; Center for Microbial Cell Biology, Leiden, The Netherlands; Technische Universität Berlin, Institute of Biotechnology, Berlin, Germany

**Keywords:** *oriC*, *Clostridioides difficile*, DnaA, P1 nuclease

## Abstract

Faithful DNA replication is crucial for viability of cells across all kingdoms of life. Targeting DNA replication is a viable strategy for inhibition of bacterial pathogens. *Clostridioides difficile* is an important enteropathogen that causes potentially fatal intestinal inflammation. Knowledge about DNA replication in this organism is limited and no data is available on the very first steps of DNA replication. Here, we use a combination of *in silico* predictions and *in vitro* experiments to demonstrate that *C. difficile* employs a bipartite origin of replication that shows DnaA-dependent melting at *oriC2,* located in the *dnaA-dnaN* intergenic region. Analysis of putative origins of replication in different clostridia suggests that the main features of the origin architecture are conserved. This study is the first to characterize aspects of the origin region of *C. difficile* and contributes to our understanding of the initiation of DNA replication in clostridia.

## 1. Introduction

*Clostridioides difficile* (formerly *Clostridium difficile*) (Lawson et al., 2016) is a Grampositive anaerobic bacterium. *C. difficile* infections (CDI) can occur in individuals with a disturbed microbiota and is one of the main causes of hospital associated diarrhea, but can also be found in the environment (Smits et al., 2016). The incidence of CDI has increased worldwide since the beginning of the century (Smits et al., 2016; Warriner et al., 2017). Consequently, the interest in the physiology of the bacterium has increased in order to understand its interaction with the host and the environment and to explore news pathways for intervention (van Eijk et al., 2017; Crobach et al., 2018).

One such pathway is the replication of the chromosome. Overall, DNA replication is a highly conserved process across different kingdoms of life (O’Donnell et al., 2013; Bleichert et al., 2017). In all bacteria, DNA replication is a tightly regulated process that occurs with high fidelity and efficiency, and is essential for cell survival. The process involves many different proteins that are required for the replication process itself, or to regulate and aid replisome assembly and activity (Katayama et al., 2010; Murray and Koh, 2014; Chodavarapu and Kaguni, 2016; Jameson and Wilkinson, 2017; Schenk et al., 2017). Replication initiation and its regulation arguably are candidates for the search of novel therapeutic targets (Fossum et al., 2008; Grimwade and Leonard, 2017; van Eijk et al., 2017).

In most bacteria, replication of the chromosome starts with the assembly of the replisome at the origin of replication *(oriC)* and proceeds bidirectionally (Chodavarapu and Kaguni, 2016). In the majority of bacteria replication is initiated by the DnaA protein, an ATPase Associated with diverse cellular Activities (AAA+ protein) that binds specific sequences in the *oriC* region. The binding of DnaA induces DNA duplex unwinding, which subsequently drives the recruitment of other proteins, such as the replicative helicase, primase and DNA polymerase III proteins {Chodavarapu, 2016 #974}. Termination of replication eventually leads to disassembly of the replication complexes (Chodavarapu and Kaguni, 2016).

In *C. difficile,* knowledge on DNA replication is limited. Though many proteins appear to be conserved between well-characterized species and *C. difficile,* only certain replication proteins have been experimentally characterized for *C. difficile* (Torti et al., 2011; Briggs et al., 2012; van Eijk et al., 2016). DNA polymerase C (PolC, CD1305) of *C. difficile* has been studied in the context of drug-discovery and appears to have a conserved primary structure similar to other low-[G+C] gram-positive organisms (Torti et al., 2011). It is inhibited *in vitro* and *in vivo* by compounds that compete for binding with dGTP (van Eijk et al., 2019; Xu et al., 2019). Helicase (CD3657), essential for DNA duplex unwinding, was found to interact in an ATP-dependent manner with a helicase loader (CD3654) and loading was proposed to occur through a ring-maker mechanism (Davey and O’Donnell, 2003; van Eijk et al., 2016). However, in contrast to helicase of the Firmicute *Bacillus subtilis, C. difficile* helicase activity is dependent on activation by the primase protein (CD1454), as has also been described for *Helicobacter pylori* (Bazin et al., 2015; van Eijk et al., 2016). *C. difficile* helicase stimulates primase activity at the trinucleotide 5’d(CTA), but not at the preferred trinucleotide 5’-d(CCC) (van Eijk et al., 2016).

DnaA of *C. difficile* has not been studied to date. Although no full-length structure has been determined for DnaA, individual domains of the DnaA protein from different organisms have been characterized (Majka et al., 1997; Zawilak et al., 2003; Erzberger et al., 2006; Zawilak-Pawlik et al., 2017). DnaA proteins generally comprise four domains (Zawilak-Pawlik et al., 2017). Domain I is involved in protein-protein interactions and is responsible for DnaA oligomerization (Weigel et al., 1999; Abe et al., 2007; Natrajan et al., 2009; Jameson et al., 2014; Kim et al., 2017; Zawilak-Pawlik et al., 2017; Martin et al., 2018; Matthews and Simmons, 2019; Nowaczyk-Cieszewska et al., 2019). Little is known about a specific function of domain II and this domain may even be absent (Erzberger et al., 2002). It is thought to be a flexible linker that promotes the proper conformation of the other DnaA domains (Abe et al., 2007; Nozaki and Ogawa, 2008). Domain III and Domain IV are responsible for the DNA binding. Domain III contains the AAA+ motif and is responsible for binding ATP, ADP and single-stranded DNA, as well as certain regulatory proteins (Kawakami et al., 2005; Cho et al., 2008; Ozaki et al., 2008; Ozaki and Katayama, 2012). Recent studies have also revealed the importance of this domain for binding phospholipids present in the bacterial membrane (Saxena et al., 2013). The C-terminal Domain IV contains a helixturn-helix motif (HTH) and is responsible for the specific binding of DnaA to so called DnaA boxes (Blaesing et al., 2000; Erzberger et al., 2002; Fujikawa et al., 2003).

DnaA boxes are typically 9-mer non-palindromic DNA sequences, and the *E. coli* DnaA box consensus sequence is TTWTNCACA (Schaper and Messer, 1995; Wolanski et al., 2014). The boxes can differ in their affinity for DnaA, and even demonstrate different dependencies on the ATP co-factor (Speck et al., 1999; Patel et al., 2017). Binding of domain IV to the DnaA boxes promotes higher-order oligomerization of DnaA, forming a filament that wraps around DNA (Erzberger et al., 2006; Ozaki et al., 2012; Scholefield and Murray, 2013). It is thought that the interaction of the DnaA filament with the DNA helix introduces a bend in the DNA (Erzberger et al., 2006; Patel et al., 2017). The resulting superhelical torsion facilitates the melting of the adjacent A+T-rich DNA Unwinding Element (DUE) (Kowalski and Eddy, 1989; Erzberger et al., 2006; Zorman et al., 2012). Upon melting, the DUE provides the entry site for the replisome proteins. Another conserved structural motif, a triplet repeat called DnaA-trio, is involved in the stabilization of the unwound region (Richardson et al., 2016; Richardson et al., 2019).

The *oriC* region has been characterized for several bacterial species. These analyses show that *oriC* regions are quite diverse in sequence, length and even chromosomal location, all of which contribute to species-specific replication initiation requirements (Zawilak-Pawlik et al., 2005; Ekundayo and Bleichert, 2019). In Firmicutes, including *C. difficile,* the genomic context of the origin regions appears to be conserved and encompasses the *rnpA-rpmH-dnaA-dnaN* genes (Ogasawara and Yoshikawa, 1992; Briggs et al., 2012).

The *oriC* region can be continuous (i.e. located at a single chromosomal locus) or bipartite (Wolanski et al., 2014). Bipartite origins where initially identified in *B. subtilis* (Moriya et al., 1988) but more recently also in *H. pylori* (Donczew et al., 2012). The separate subregions of the bipartite origin, *oriC1* and *oriC2,* are usually separated by the *dnaA* gene. Both *oriC1* and *oriC2* contain clusters of DnaA boxes, and one of the regions contains the major DUE region. The DnaA protein binds to both subregions and places them in close proximity to each other, consequently looping out the *dnaA* gene (Krause et al., 1997; Donczew et al., 2012). In *H. pylori,* DnaA domain I and II are important for maintaining the interactions between both *oriC* regions (Nowaczyk- Cieszewska et al., 2019).

In this study, we identified the putative *oriC* of *C. difficile* through *in silico* analysis and demonstrate DnaA-dependent unwinding of the *oriC2* region *in vitro.* A clear conservation of the origin of replication organization is observed throughout the clostridia. The present study contributes to our understanding of clostridial DNA replication initiation in general, and replication initiation of *C. difficile* specifically.

## 2. Materials and Methods

### 2.1 Sequence alignments and structure modelling

Multiple sequence alignment of amino acid sequences was performed with Protein BLAST (blastP suite, https://blast.ncbi.nlm.nih.gov/Blast.cgi) for individual alignment scores and the PRALINE program (http://www.ibi.vu.nl/programs/pralinewww/) (Bawono and Heringa, 2014) for multiple sequence alignment. Sequences were retrieved from the NCBI Reference Sequences. DnaA protein sequences from *C. difficile* 630Δ*erm* (CEJ96502.1), *C. acetobutylicum* DSM 1731 (AEI33799.1), *Bacillus subtilis* 168 (NP_387882.1), *Escherichia coli* K-12 (AMH32311.1), *Streptomyces coelicolor* A3(2) (TYP16779.1), *Mycobacterium tuberculosis* RGTB327 (AFE14996.1), *Helicobacter pylori J99* (Q9ZJ96.1) and *Aquifex aeolicus* (WP_010880157.1) were selected for alignment. Alignment was visualized in JalView version 2.11, with coloring by percentage identity.

Secondary structure prediction and homology modelling were performed using Phyre2 (http://www.sbg.bio.ic.ac.uk/phyre2) (Kelley et al., 2015) using the intensive default settings. Phyre2 modelling of *C. difficile 630Δerm* DnaA (CEJ96502.1) was performed with 3 templates from *A. aeolicus* (PDB 2HCB, chain C), *B. subtilis* (PDB 4TPS, chain D) and *E. coli* (PDB 2E0G, chain A) and 21 residues were modelled *ab initio.* 95% of the residues were modelled with >90% confidence. Graphical representation was performed with the PyMOL Molecular Graphics System, Version 1.76.6. Schrödinger, LLC.

### 2.2 Prediction of the *C. difficile oriC*

To identify the *oriC* region of *C. difficile* the genome sequence of *C. difficile 630Δerm* (GenBank accession no. LN614756.1) was analyzed through different software in a stepwise procedure (Mackiewicz et al., 2004).

The GenSkew Java Application (http://genskew.csb.univie.ac.at/) was used with default settings for the analysis of the normal and the cumulative skew of two selectable nucleotides of the genomic nucleotide sequence ([G – C]/[G + C]). Calculations where performed with a window size of 4293 bp and a step size of 4293 bp. The inflection values of the cumulative GC skew plot are indicative of the chromosomal origin (*oriC*) and terminus of replication (*ter*).

Prediction of superhelicity-dependent helically unstable DNA stretches (SIDDs) was performed in the vicinity of the inflection point of the GC-skew plot, in 2.0 kb fragments comprising intergenic regions from nucleotide position 4291795 to 745 *(oriC1)* and 466 to 2465 *(oriC2)* of the *C. difficile 630Δerm* chromosome. Prediction of the SIDDs in the different clostridia (Table 1) was performed in the vicinity of the inflection points of the GC-plot retrieved from DoriC 10.0 database (http://tubic.tju.edu.cn/doric/public/index.php) (Luo and Gao, 2019), in 2.0 kb fragments comprising intergenic regions summarized in Table 1. The SIST program (https://bitbucket.org/benhamlab/sist_codes/src/master/) (Zhabinskaya et al., 2015) was used to predicted free energies G(x) by running the melting transition algorithm only (SIDD) with default values (copolymeric energetics; default: σ =–0.06; T = 37°C; x= 0.01M) and with superhelical density σ = −0.04.

**Table 1.** Clostridia intergenic regions used for SIDD analysis.

| Clostridia (GenBank accession no.) | *oriC1*<sup>*1</sup><br>*DoriC ID*<sup>*2</sup> | *oriC2*<br>*DoriC ID*<sup>*</sup> |
| --- | --- | --- |
| *C. difficile* R20291 (NC_013316.1) | 4189900 to 561<br>ORI93010593 | 780 to 2780<br>ORI93010592 |
| *C. botulinum* A Hall (NC_009698.1) | 3759361 to 800<br>ORI92010336 | 510 to 2510<br>ORI92010335 |
| *C. sordelli* AM370 (NZ_CP014150 | 3549121 to 662<br>ORI97012279 | 561 to 2561<br>ORI97012278 |
| *C. acetobutylicum* DSM 1731 (NC_015687.1) | 3941422 to 961<br>ORI94010884 | 1040 to 3040<br>ORI94010883 |
| *C. perfringens* str.13 (NC_003366.1) | 3030241 to 810<br>ORI10010054 | 881 to 2881<br>ORI10010053 |
| *C. tetani* E88 (NC_004557.1) | 52001 to 54000<br>ORI10010089 | 50081 to 52081<br>ORI10010088 |

<sup>\*1</sup> 2.0 kb fragments selected for SIDD analysis comprising the intergenic regions
<sup>\*2</sup> DoriC 10.0 intergenic regions from http://tubic.tju.edu.cn/doric/public/index.php

We performed the identification of the DnaA box clusters by search of the motif TTWTNCACA with one mismatch (Supplementary information) in the leading strand on a 4432 bp sequence between the nucleotide position 4291488 to 2870 of the *C. difficile 630Δerm* chromosome, using Pattern Locator (https://www.cmbl.uga.edu//downloads/programs/Pattern_Locator/patloc.c) (Mrazek and Xie, 2006). Identification of the DnaA boxes in the different clostridia (Table 1) was performed with the same pattern motif in the leading strand of the intergenic regions summarized on Table 1.

DnaA-trio sequences and ribosomal binding sites where manually predicted based on Richardson et all. (Richardson et al., 2016) and on Vellanoweth and Rabinowitz (Vellanoweth and Rabinowitz, 1992), respectively.

All output data was obtained as raw text files and further processed with Prism 8.3.1 (GraphPad, Inc, La Jolla, CA) and CorelDRAW X7 (Corel).

### 2.3 Strains and growth conditions

*E. coli* strains were grown aerobically at 37°C in lysogeny broth (LB, Affymetrix) supplemented with 15 μg/mL chloramphenicol or 50 μg/mL kanamycin when required. *E. coli* strain DH5α (Table 2) for DnaA containing plasmid and *E. coli* MC1061 strain (Table 2) was used to maintain the *oriC* containing plasmids. *E. coli* MS3898 strain, kindly provided by Alan Grossman (MIT, Cambridge, USA) (Table 2) was used for recombinant DnaA expression. *E. coli* transformation was performed using standard procedures (Sambrook et al., 1989). The growth was followed by monitoring the optical density at 600 nm (OD_600_).

**Table 2.** *E. coli* strains used in this study.

| Name | Relevant Genotype/Phenotype* | Origin |
| --- | --- | --- |
| DH5α | F– endA1 glnV44 thi-1 recA1 relA1 gyrA96 deoR nupG purB20 ϕ80dlacZΔM15 Δ(lacZYA-argF)U169, hsdR17(rK–mK+), λ– | Laboratory collection |
| MC1061 | str. K-12 F– λ– Δ(ara-leu)7697 [araD139]B/r Δ(codB-lacI)3 galK16 galE15 e14– mcrA0 relA1 rpsL150(StrR) spoT1 mcrB1 hsdR2(r–m+) | Laboratory Collection |
| CYB1002 | ΔdnaA zia::pKN500(miniR1) asnB32 relA1 spoT1 thi-1 ilv192 mad1 recA1 λimm434 F- pBB42 (lacI; TetR) | Grossman lab |

### 2.4 Construction of the plasmids

For overexpression of DnaA, the *dnaA* nucleotide sequence (CEJ96502.1) from *C. difficile 630Δerm* (GenBank accession no. LN614756.1) was amplified by PCR from *C. difficile 630Δerm* genomic DNA using primers oEVE-7 and oEVE-21 (Table 3). The PCR product was subsequently digested with NcoI and BglII. The vector pAV13 (Smits et al., 2011) (Table 4), containing *B. subtilis dnaA* cloned in pQE60 (Qiagen) was kindly provided by Alan Grossman (MIT, Cambridge, USA) and was digested with the same enzymes and ligated to the digested fragment to yield vector pEVE40 (Table 4).

**Table 3.**
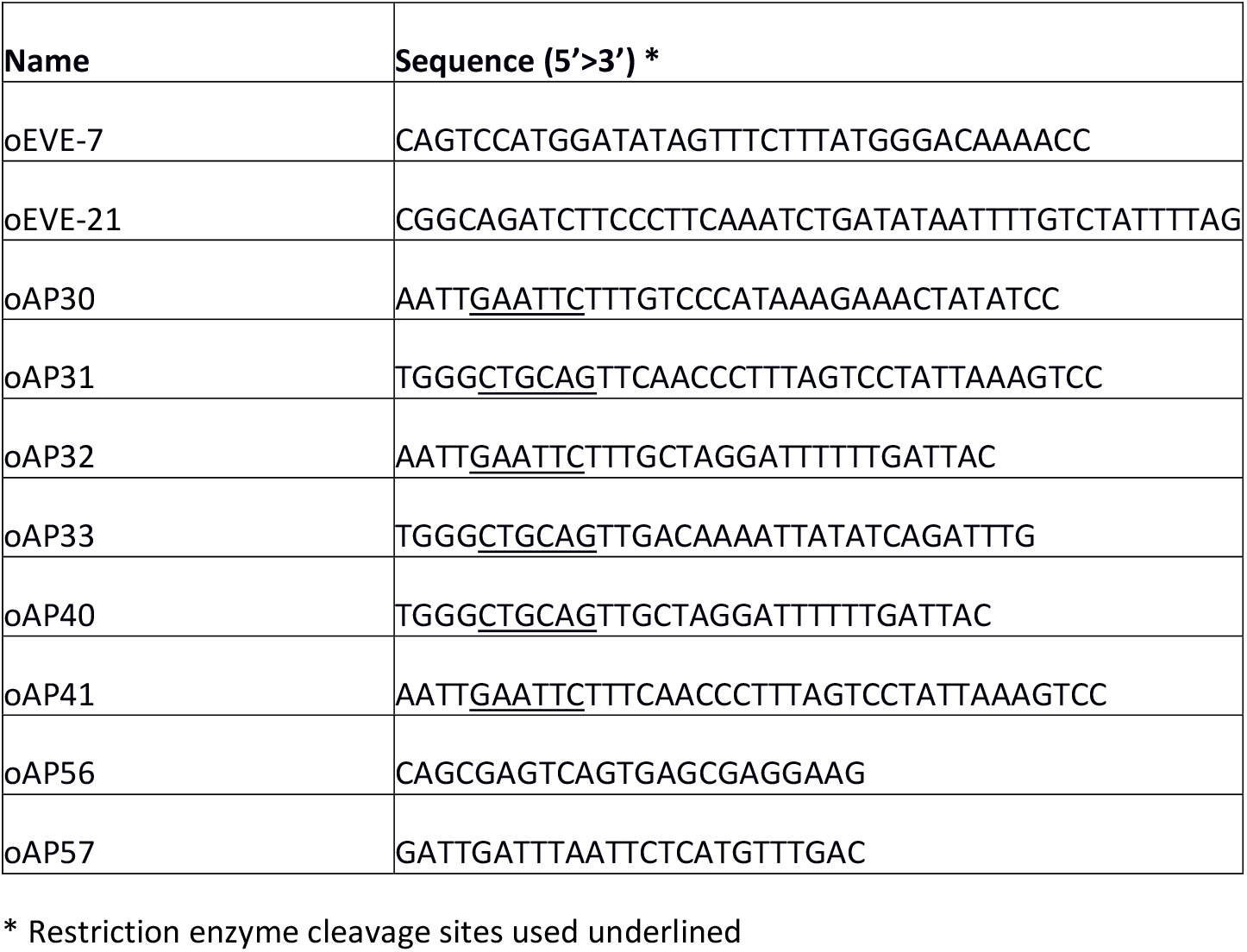
Oligonucleotides used in this study.

**Table 4.** Plasmids used in this study.

| Name | Relevant features* | Source/Reference |
| --- | --- | --- |
| pAV13 | lacI<sup>q</sup>, P<sub>T5</sub> expression vector; *km* | (Smits, Merrikh et al. 2011) |
| pEVE40 | P<sub>T5</sub> - DnaA-6xHis; *km* | This study |
| pori1ori2 | *H. pylori oriC1oriC2; amp* | (Donczew, Weigel et al. 2012) |
| pAP76 | *C. difficile oriC2; amp* | This study |
| pAP83 | *C. difficile oriC1; amp* | This study |
| pAP205 | *C. difficile oriC1oriC2; amp* | This study |

To construct a plasmid carrying the complete predicted *oriC,* the predicted *oriC* region (nucleotide 4292150 to 1593 from *C. difficile* 630 GenBank accession no. LN614756.1) was amplified by PCR from *C. difficile 630Δerm* genomic DNA using primers oAP40 and oAP41 (Table 3). The PCR product was subsequently digested with EcoRI and PstI and ligated into pori1ori2 (Table 4), kindly provided by Anna Zawilak-Pawlik (Hirszfeld Institute of Immunology and Experimental Therapy, PAS, Wroclaw, Poland), that was digested with the same enzymes, to yield vector pAP205 (Table 4).

For the cloning of the predicted *oriC1* region (nucleotide 4292150 to 24 of *C. difficile 630Δerm* genomic DNA) the primer set oAP30/oAP31 (Table 3) was used. The amplified fragment was digested with EcoRI and PstI and inserted onto pori1ori2 (Table 4) digested with same enzymes, yielding vector pAP83 (Table 4). For the cloning of the predicted *oriC2* region (nucleotide 1291 to the 1593 of *C. difficile 630Δerm* genomic DNA) the primer set oAP32/oAP33 (Table 3) was used. The amplified fragment was digested with EcoRI and PstI and inserted onto pori1ori2 (Table 4) digested with same enzymes, yielding vector pAP76 (Table 4).

All DNA sequences introduced into the cloning vectors were verified by Sanger sequencing. For *oriC* containing vectors primers oAP56 and oAP57 (Table 3) were used for sequencing.

### 2.5 Overproduction and purification of DnaA-6xHis

Overexpression of DnaA-6xHis was carried out in *E. coli* strain CYB1002 (Table 2), harbouring the expression plasmid pEVE40 (Table 4). Cells were grown in 800 mL LB and induced with 1mM isopropyl-β-D-1-thiogalactopyranoside (IPTG) at an OD_600_ of 0.6 for 3 hours. The cells were collected by centrifugation at 4°C and stored at −80°C. Cells were resuspended in Binding buffer (1X Phosphate buffer pH7.4, 10 mM Imidazol, 10% glycerol) lysed by French Press and collected in phenylmethylsulfonyl fluoride (PMSF) at 0.1 mM (end concentration). Separation of the soluble fraction was performed by centrifugation at 13000xg at 4°C for 20 min. Purification of the protein from the soluble fraction was done in Binding buffer on a 1 mL HisTrap Column (GE Healthcare) according to manufacturer’s instructions. Elution was performed with Binding buffer in stepwise increasing concentrations of imidazole (20, 60, 100, 300 and 500 mM). DnaA- 6xHis was mainly eluted at concentration of imidazole equal to or greater than 300mM.

Fractions containing the DnaA-6xHis protein were pooled together and applied to Amicon Ultra Centrifugal Filters with 30 kDa cutoff (Millipore). Buffer was exchanged to Buffer A (25 mM HEPES-KOH pH 7.5, 100 mM K-glutamate, 5 mM Mg-acetate, 10% glycerol). The concentrated DnaA protein was subjected to size exclusion chromatography on an Äkta pure instrument (GE Healthcare). 200 μL of concentrated DnaA-6xHis was applied to a Superdex 200 Increase 10/30 column (GE Healthcare) in buffer A at a flow rate of 0.5 ml min^-1^. UV detection was done at 280 nm. The column was calibrated with a mixture of proteins of known molecular weights (Mw): thyroglobulin (669 kDa), Apoferritin (443 kDa), β-amylase (200 kDa), Albumin (66 kDa) and Carbonic anhydrase (29 kDa). Eluted fractions containing DnaA-6xHis of the expected molecular weight (51 kDa) were quantified and visualized by Coomassie. Pure fractions were aliquoted and stored at −80°C for further experiments.

### 2.6 Immunoblotting and detection

For immunoblotting, proteins were separated on a 12% SDS-PAGE gel and transferred onto nitrocellulose membranes (Amersham), according to the manufacturer’s instructions. The membranes were probed in PBST (PBS pH 7,4, 0,05% (v/v) Tween-20) with the mouse anti-his antibody (1:3000, Invitrogen) and the respective secondary antibody goat anti-mouse-HRP (1:3000, DAKO) were used. The membranes were visualized using the chemiluminescence detection kit Clarity ECL Western Blotting Substrates (Bio-Rad) in an Alliance Q9 Advanced machine (Uvitec).

### 2.7 P1 nuclease Assay

For the P1 nuclease assay, 100 ng pAP205 plasmid was incubated with increasing concentrations of DnaA-6xHis (0.14, 0.54, 1 and 6.3 μM), when required, in P1 buffer (25mM Hepes-KOH (pH 7.6), 12% (v/v) glycerol, 1mM CaCl_2_, 0.2mM EDTA, 5mM ATP, 0.1 mg/ml BSA), at 30°C for 12 min. 0.75 unit of P1 nuclease (Sigma), resuspended in 0.01 M sodium acetate (pH 7.6) was added to the reaction and incubated at 30°C for 5 min. 220 μl of buffer PB (Qiagen) was added and the fragments purified with the miniElute PCR Purification Kit (Qiagen), according to manufacturer’s instructions. Digestion with BglII, NotI or ScaI (NEB) of the purified fragments was performed according to manufacturer’s instructions for 1 hour at 37°C. Digested samples were resolved on 1% agarose gels in 0.5xTAE (40 mM Tris, 20 mM CH_3_COOH, 1 mM EDTA PH 8.0) and stained with 0.01 mg/mL ethidium bromide solution afterwards. Visualization of the gels was performed on the Alliance Q9 Advanced machine (Uvitec). Images were processed in CorelDraw X7 software.

## 3. Results

### 3.1 *C. difficile* DnaA protein

*C. difficile 630Δerm* encodes a homolog of the bacterial replication initiator protein DnaA (GenBank: CEJ96502.1; CD630DERM_00010). Alignment of the full-length *C. difficile* DnaA amino acid sequence with selected DnaA homologs from other organisms demonstrates a sequence identity of 35% to 67%, with an even higher similarity (57% to 83%, Fig. 1A). *C. difficile* DnaA displays a greater sequence identity between the low-[G+C] Firmicutes (> 60%). When compared with the extensively studied DnaA proteins from *E. coli* and *B. subtilis,* the full-length protein has 43% and 62% identity, and a similarity of 63% and 78%, respectively (Fig. 1A).

**Figure 1.**
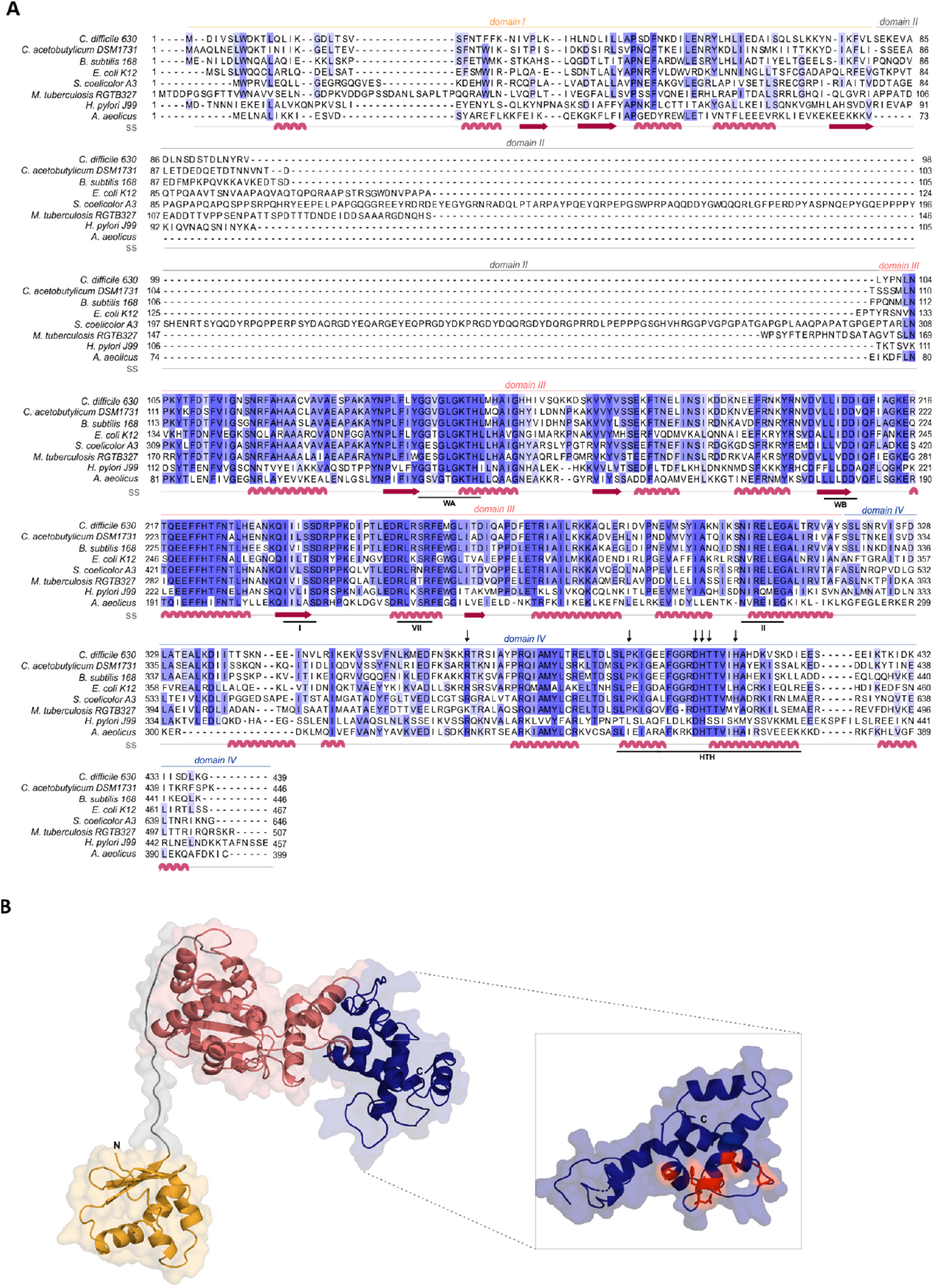
*C. difficile* DnaA DNA binding domain is conserved. **A)** Multiple sequence alignment (PRALINE) of *C. difficile* DnaA with homologous proteins retrieved from GenBank. The aminoacid sequences from *C. difficile* 630Δerm (CEJ96502.1), *C. acetobutylicum* DSM 1731 (AEI33799.1), *B. subtilis* 168 (NP_387882.1), *E. coli* K-12 (AMH32311.1), *S. coelicolor* A3(2) (TYP16779.1), *M. tuberculosis* RGTB327 (AFE14996.1), *H. pylori* J99 (Q9ZJ96.1) and *Aquifex aeolicus* (WP_010880157.1) were used. Residues are colored according to sequence identity conservation highlighted with blue shading (dark blue more conserved), performed in JalView. Secondary structure prediction (ss) is indicated, according to Phyre2 modelled structure. DnaA domains are represented, with the conserved AAA+ ATPase fold motifs Walker A, Walker B, VII box, sensor I and sensor II highlighted (WA, WB, I, VII and II motifs), as well as the domain IV helix-turn-helix (HTH). Residues involved in the base-specific recognition are identified with an arrow. **B)** Structural model of *C. difficile* DnaA determined by Phyre2. Domains are colored as in alignment. Both the N-terminus and the C-terminus are indicated in the figure. The DnaA domain IV is enhanced (inset) with the DnaA-box binding specific residues represented in red sticks.

To assess the structural properties of *C. difficile* DnaA, we predicted the secondary structure and generated a model of the protein using Phyre2 (Kelley et al., 2015) (Fig. 1B). The predicted DnaA model is based on three DnaA structures from different organisms: *A. aeolicus* (residues 101 to 318 and 334 to 437)(Erzberger et al., 2006) for domain III and IV, and *B. subtilis* (residues 2 to 79) (Jameson et al., 2014) and *E. coli* (residues 5 to 97) (Abe et al., 2007) for domain I and II.

Domain I of DnaA mediates interactions with a diverse set of regulators, and is involved in DnaA oligomerization (Zawilak-Pawlik et al., 2017; Nowaczyk-Cieszewska et al., 2019). We observe limited homology of *C. difficile* DnaA domain I with the equivalent domain of the selected organisms (Fig. 1A), although the overall fold is clearly conserved (Fig. 1B). Nevertheless, some residues (P45, F48) appear to be conserved in most of the selected organisms (Fig. 1A).

Domain II is a flexible linker that is possibly involved in aiding the proper conformation of the DnaA domains, and thus requires a minimal length for DnaA function *in vivo* (Nozaki and Ogawa, 2008). No clear sequence similarity is observed on domain II and modelling of the *C. difficile* DnaA protein suggests a putative disordered nature of this domain (Fig. 1).

Domain III is responsible for binding to the co-factors ATP and ADP, and in conjunction with domain IV essential for DNA binding (Kawakami et al., 2005; Ozaki et al., 2008; Ozaki and Katayama, 2012). Within domain III we readily identified the Walker A and Walker B motifs (WA and WB in Fig. 1A) of the AAA+ fold (residues 135-317), crucial for binding and hydrolyzing ATP. This domain is highly conserved among all the selected organisms (Fig. 1A) and comprises a structural center of β-sheets (Fig. 1B, pink domain). Other features of the AAA+ ATPase fold are present and conserved between the organisms, such as the sensor I and sensor II motifs required for the nucleotide binding (I and II, Fig. 1A). The arginine finger motif (the equivalent of R285 of *E.coli* DnaA in the VII box), important for the ATP dependent activation of DnaA (Kawakami et al., 2005), is conserved in *C. difficile* DnaA as well (R256 in motif box VII; Fig. 1A).

The C-terminal domain IV of the DnaA protein (residues 317 to 439, Fig. 1A), contains the HTH motif required for the specific binding to DnaA-boxes (Erzberger et al., 2002; Zawilak et al., 2003). Previous studies identified several residues involved in specific interactions with the DnaA boxes, that bind through hydrogen bonds and van der Waals contacts with thymines present in the DNA sequence (Blaesing et al., 2000; Fujikawa et al., 2003; Tsodikov and Biswas, 2011). The residues are conserved among all Firmicutes and *E. coli,* including the residues R371 (position R399 in *E. coli),* P395 (P423), D405 (D433), H406 (H434), T407 (T435), and H411 (H439), (Fig. 1B inset, red residues) (Fujikawa et al., 2003). Structural modeling of *C. difficile* DnaA predicts these residues to be exposed, providing an interface for DNA binding (Fig. 1B). Several residues were found to be involved in non-specific interactions with the phosphate backbone of the DNA, including some of the residues that confer the specificity (Fujikawa et al., 2003; Tsodikov and Biswas, 2011). These contacting residues appear less conserved between the selected organisms (Fig. 1A. Nevertheless, the residues for specific base recognition are conserved between the Firmicutes and *E. coli,* suggesting that *C. difficile* DnaA is likely to recognize the consensus DnaA box TTWTNCACA (Schaper and Messer, 1995).

### 3.2 Expression and purification of DnaA-6xHis

To allow for *in vitro* characterization of DnaA activity, we recombinantly expressed the *C. difficile* DnaA with a C-terminal 6xHis-tag in *E. coli* cells. To prevent the copurification of *C. difficile* DnaA with host DnaA protein, *E. coli* strain CYB1002 was used (a kind gift of A.D. Grossman). This strain is a derivative of *E. coli* MS3898, that lacks the *dnaA* gene and replicates in a DnaA-independent fashion (Sutton and Kaguni, 1997). Induction of the DnaA-6xHis protein was confirmed by Coomassie staining and immunoblotting with anti-his antibody at the expected molecular weight of 51 kDa (Fig. 2A, red arrow). Upon overexpression of DnaA-6xHis, smaller fragments were observed, which accumulated with a prolonged time of expression (Fig. 2A), most likely corresponding to proteolytic fragments of the DnaA-6xHis protein.

**Figure 2.**
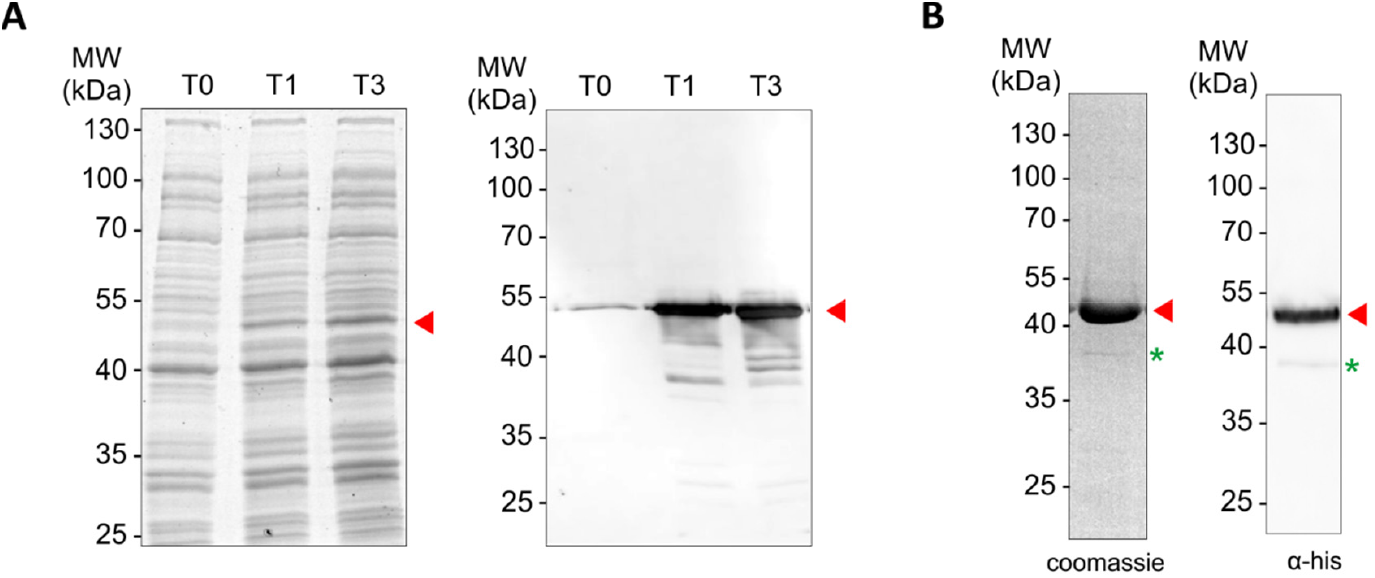
Expression and purification of *C. difficile* DnaA protein. **A)** *E. coli* expressing DnaA-6xHis cells were induced with 1 mM IPTG. Optical density-normalized samples before induction (T0), after 1 hour of induction (T1) and 3 hours of induction (T3) were resolved by 12% SDS-PAGE and immunoblotted with anti-his antibody. Induced DnaA is observed with the approximate molecular weight of 51 kDa (red arrow). Possible breakdown product is observed (blue arrow). **B)** Confirmation of size-exclusion fraction containing the *C. difficile* DnaA-6xHis and further used for analysis after protein purification resolved by 12% SDS-PAGE (Coomassie staining) and immunoblotted with anti-his antibody. DnaA-6xHis is observed with the approximate molecular weight of ~51 kDa (red arrow). Possible minor breakdown products are observed (green asterisk).

Purification of the recombinant DnaA-6xHis showed a clear band at the expected size when eluted at 300 mM imidazole concentration, but several lower molecular size bands were observed (Fig. S1). Therefore, the eluted fractions where further purified with size exclusion chromatography (SEC). This yielded a single product at the expected molecular weight of DnaA-6xHis, and its identity was confirmed by westernblot with anti-his antibody (Fig. 2B, red arrow). A minor band of lower molecular weight (approximately 38 kDa, <1% of total protein) was observed (Fig. 2B, green asterisk), which may reflect some instability of the N-terminus of the DnaA-6xHis protein, as it appears to have retained the C-terminal 6xHis tag.

### 3.3 *In silico* prediction of the *oriC* region

To identify the *oriC* region and the elements that are part of it (DUE, DnaA-trio and DnaA boxes) we performed different prediction approaches in a stepwise procedure, as initially described (Mackiewicz et al., 2004).

We first analyzed the DNA asymmetry of the genome of *C. difficile 630Δerm* (GenBank accession no. LN614756.1) (van Eijk et al., 2015), by plotting the normalized difference of the complementary nucleotides (GC-skew plot) (Necsulea and Lobry, 2007). *C. difficile 630Δerm* has a circular genome of 4293049 bp and an average G+C content of 29.1%. We used the GenSkew Java Application (http://genskew.csb.univie.ac.at/) for determining the chromosomal asymmetry. Asymmetry changes in a GC-skew plot can be used to predict the origin of replication region and the terminus region of bacterial genomes. Based on this analysis, the origin is predicted at approximately position 1 of the chromosome. The terminus location is predicted at approximately 2.18 Mbp from the origin region (Fig. 3A). These results were confirmed when artificially reassigning the starting position of the chromosomal assembly (data not shown). The gene organization in the putative origin region is *rnpA-rpmH-dnaA-dnaN* (position 4291488 to 2870, Fig. 3B), identical to the origin of *B. subtilis* (Ogasawara et al., 1985; Briggs et al., 2012), and therefore encompasses the *dnaA* gene (CD630DERM_00010, Fig. 3B) (Ogasawara et al., 1985; Briggs et al., 2012).

**Figure 3.**
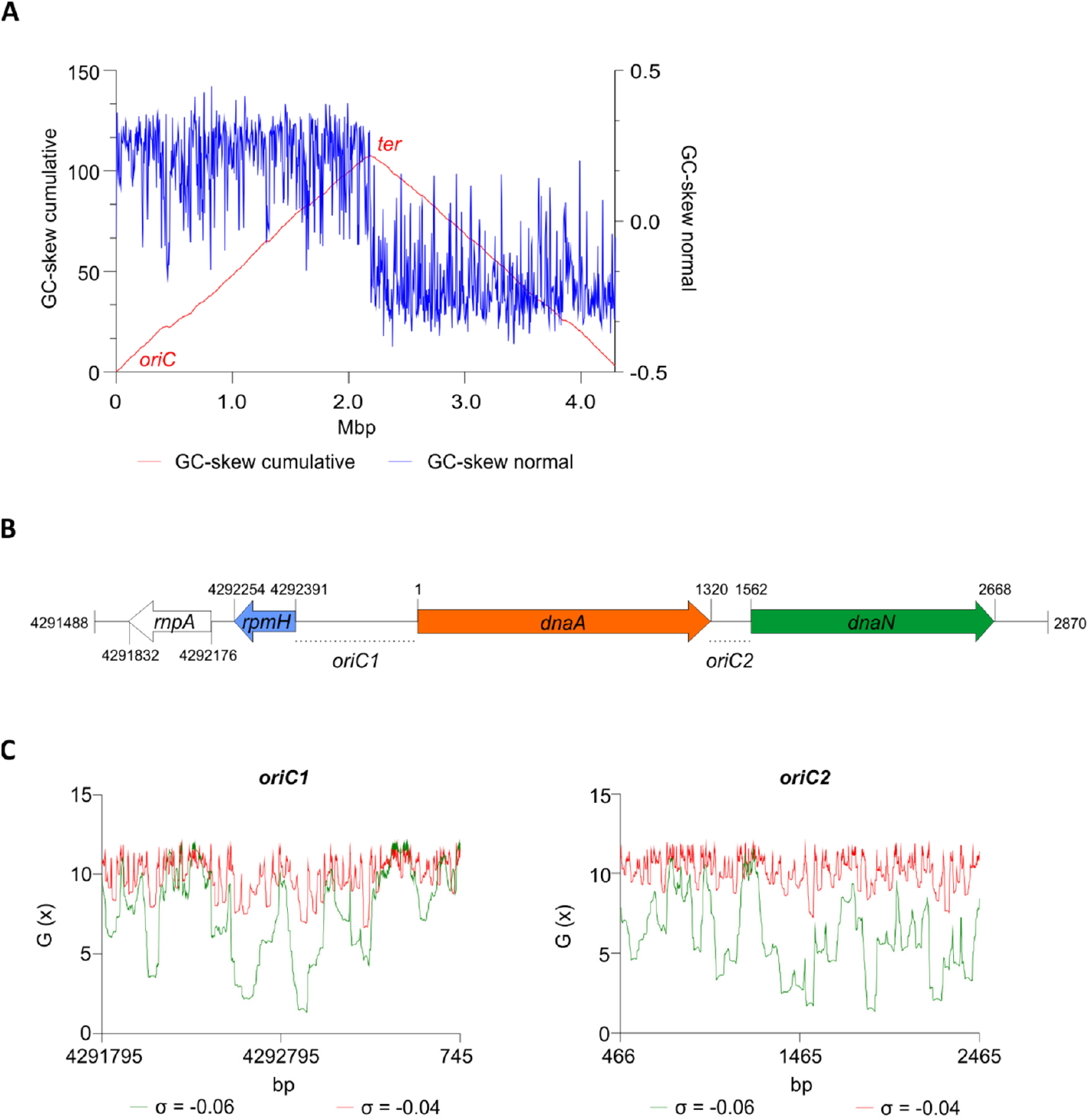
Prediction of the *C. difficile* origin of replication. **A)** GC skew analysis of the *C. difficile* 630Δerm (LN614756.1) genome sequence. Normal GC skew analysis ([G – C]/[G + C]) performed on leading strand (blue line) and respective cumulative GC skew plot (red line). Calculations where performed with a window size of 4293 bp and a step size of 4293 bp. The *origin (oriC) and terminus (ter) regions* are indicated. **B)** Representation of the predicted origin region and genomic context (from residues at position 4291488 to 2870 of the *C. difficile* 630 Δerm chromosome). The *rnpA, rpmH* (blue arrow), *dnaA* (orange arrow) and *dnaN* (green arrow) genes are indicated. Putative origins in intergenic regions are represented *oriC1 (rpmH-dnaA)* and *oriC2 (dnaA-dnaN).* **C)** SIDD analysis of 2.0 kb fragments comprising *oriC1* (nucleotide 4291795 to 745) and *oriC2* (nucleotide 466 to 2465). Predicted free energies G(x) for duplex destabilization at a superhelical density of σ = −0.06 (green) or σ = −0.04 (red).

We next used the SIST program (Zhabinskaya et al., 2015) to localize putative DUEs in the intergenic regions in the chromosomal region predicted to contain the *oriC.* Hereafter we refer to these regions as *oriC1* (in the intergenic region of *rpmH-dnaA)* and *oriC2* (in the intergenic region *dnaA-dnaN),* in line with nomenclature in other organisms (Ogasawara et al., 1985; Donczew et al., 2012) (Fig. 3B). SIST identifies helically unstable AT-rich DNA stretches (Stress-Induced Duplex Destabilization regions; SIDDs) (Donczew et al., 2012; Zhabinskaya et al., 2015). In regions with a lower free energy (G_(x)_ < y kcal/mol) the double-stranded helix has a high probability to become single-stranded DNA. With increasing negative superhelicity (σ = −0.06, Fig. 3C, green line) regions of both *oriC1* and *oriC2* become single stranded DNA (G_(x)_ <2 kcal/mol). At low negative superhelicity (σ = −0.04, Fig. 3C, red line) short stretches of DNA of approximately 27 bp were identified with a significantly lower free energy. These regions with lower free energy at a negative superhelicity of −0.04 and −0.06 are potential DUE sites. The nucleotide sequence of the possible unwinding elements identified are represented in detail in Fig. 4 (grey boxes).

**Figure 4.**
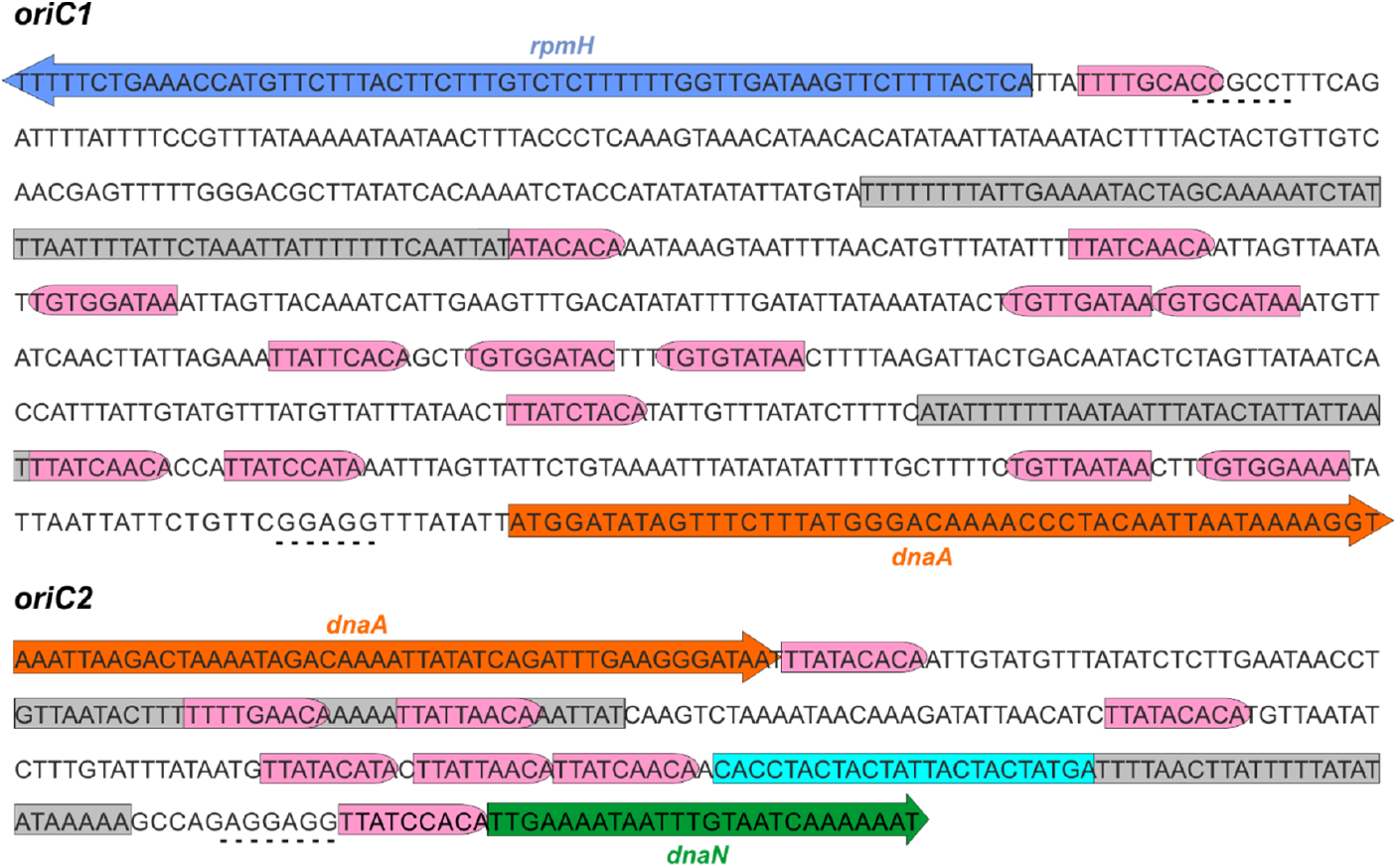
Identification of the *C. difficile oriC* region. Nucleotide sequence of the *oriC1* region (nucleotide 4292328 to 48 of the *C. difficile* 630Δerm LN614756.1 genome sequence) and *oriC2* region (nucleotide 1274 to 1587). Identification of the possible unwinding AT-rich regions previously identified in the SIDD analysis (grey boxes). The putative DnaA boxes found are represented (pink boxes) and orientation in the leading (right) and lagging strand (left) are shown. Possible DnaA-trio sequence are denoted (light blue boxes). Coding sequence of the genes *rpmH* (blue arrow), *dnaA* (orange arrow) and *dnaN* (green arrow) and respective putative ribosome binding sites (dashed line) are indicated. Pattern identification is described in Material and Methods.

We then performed the identification of DnaA box clusters through a search of the consensus DnaA box TTWTNCACA containing up to one mismatch, using Pattern Locator (Mrazek and Xie, 2006). 22 putative DnaA boxes where identified in both the leading and lagging strain in the predicted *C. difficile oriC* regions (Fig. 4, pink boxes), 14 in the *oriC1* region and 8 in the *oriC2* region. Both the consensus DnaA box TTWTNCACA and variant boxes are found. A cluster of DnaA boxes was proposed to contain at least three boxes with an average distance lower than 100 bp in between (Mackiewicz et al., 2004). At least one such cluster can be found in each origin region (Fig. 4).

We also manually identified the putative ribosomal binding sites for the annotated genes (Fig. 4, dashed line).

Finally, we manually predicted DnaA-trio sequences (3’-[G/A]A[T/A]_n>3_-5’ preceded by a GC-cluster) in the predicted *oriC* regions, as this motif is required for successful replication in both *E. coli* and *B. subtilis* (Richardson et al., 2016; Katayama et al., 2017). We identified a clear DnaA-trio in the lagging strand upstream of a predicted DUE region in the *oriC2* region, with the nucleotide sequence 5’- CACCTACTACTATTACTACTATGA-3’ (Fig. 4, light blue box), but no clear DnaA-trio was identified in the *oriC1* region.

From all the observations, we anticipate that a bipartite origin is located in the *dnaA* chromosomal region of *C. difficile* with unwinding occurring downstream of *dnaA,* at the *oriC2* region.

### 3.4 DnaA-dependent unwinding

To localize DnaA-dependent unwinding of *oriC,* we used the purified *C. difficile* DnaA- 6xHis protein and the predicted *oriC* sequence, to perform P1 nuclease assays as previously described (Sekimizu et al., 1988; Donczew et al., 2012). Localized melting resulting from DnaA activity exposes ssDNA to the action of the ssDNA-specific P1 nuclease. After incubation of a vector containing the *oriC* fragment with DnaA protein and cleavage by the P1 nuclease, the vector is purified and digested with different endonucleases to map the location of the unwound region.

We constructed vectors, based on pori1ori2 (Donczew et al., 2012), harboring *C. difficile oriC1* (pAP76) or *oriC2* (pAP83) individually, as well as the complete *oriC* region (pAP205) (Fig. 5A and S2A). For a more accurate determination of the unwound region, the vectors were subjected to digestion by three different restriction enzymes (BglII, NotI, ScaI), resulting in different restriction patterns. Limited spontaneous unwinding of the plasmid was observed in the *C. difficile oriC*-containing vectors (Fig. 5A and S2B). No DnaA-dependent change in restriction pattern was observed when using the single *oriC* regions (Fig. S2B), suggesting *oriC1* and *oriC2* individually lack the requirements for DnaA-dependent unwinding.

**Figure 5.**
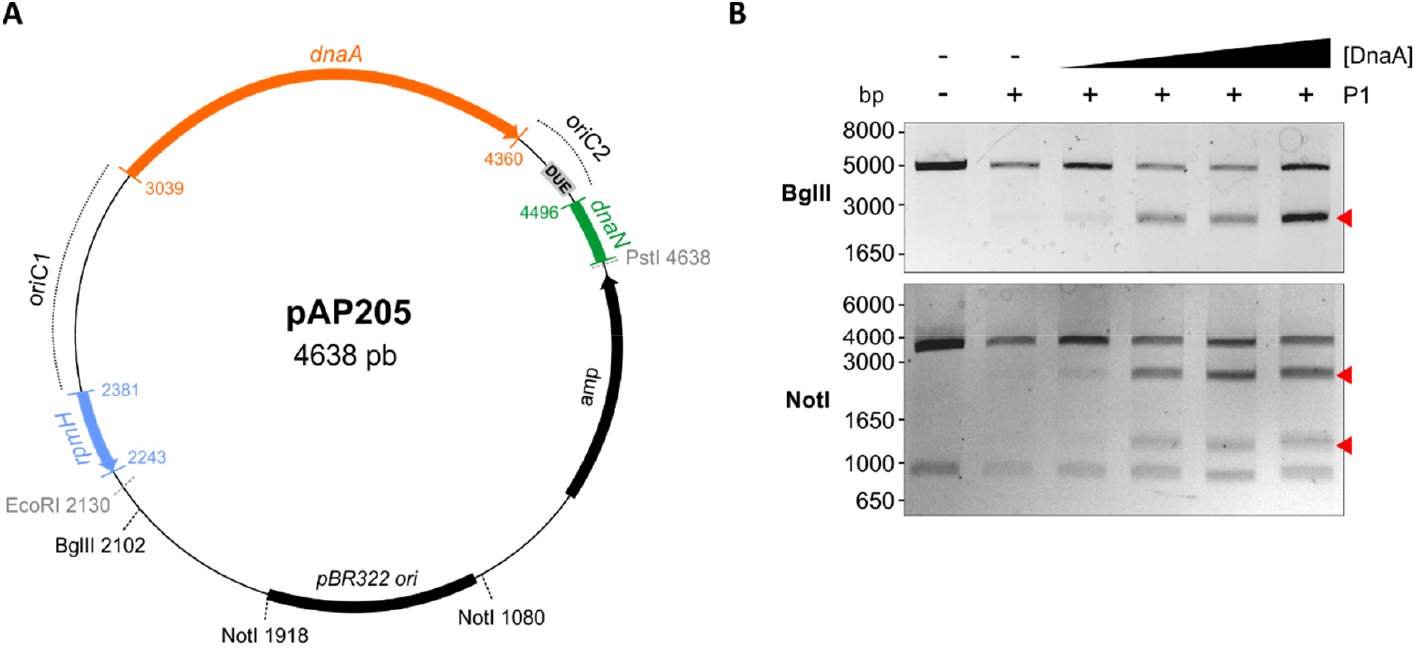
Identification of the unwinding region in *C. difficile oriC.* **A)** Representation of the *oriC1oriC2* containing vector pAP205 used in the P1 nuclease assay. The predicted *oriC1* and *oriC2* regions (dotted lines) and included genes are represented, *rpmH* (blue), *dnaA* (orange), and *dnaN* (green). The *bla* gene, the pBR322 plasmid origin of replication and the positions of used restriction sites are marked. The unwinding region (DUE) is denoted in a grey circle. **B)** P1 nuclease assay of the *or/C1oriC2*-containing vector pAP205. Digestion of the vector (lane 1) with different restriction enzymes BglII (upper panel), NotI (middle panel) and ScaI (bottom panel). Treatment of the fragments with P1 nuclease only (lane 2) and incubated with increasing amounts of *C. difficile* DnaA protein (lanes 3-6). The DNA fragments were separated in a 1% Agarose gel and analyzed with ethidium bromide staining. Resulting fragments of the DnaA-dependent unwinding are indicated with a red arrow (see results for details).

We did observe a DnaA-dependent change in digestion patterns for the *oriC1oriC2-* containing vector pAP205 (Fig 5). Digestion of this vector with BglII in the absence of DnaA-6xHis and P1 nuclease resulted in a linear DNA fragment (4638 bp) due the presence of a unique BglII restriction site (Fig. 5B, first lane, upper panel). The addition of P1 nuclease leads to the appearance of a faint band between 1650 and 3000 bp (Fig. 5B), consistent with previous observations that the presence of a plasmid DUE can result in low-level spontaneous unwinding due to the inherent instability of these AT-rich regions (Jaworski et al., 2016). Upon the addition of the DnaA-6xHis protein the observed band becomes more intense, suggesting a strong increase in unwinding (Fig. 5B, upper panel, red arrow).

Digestion of pAP205 with NotI in the absence of DnaA-6xHis and P1 nuclease results in fragments of 3804 and 842 bp, due to two NotI recognition sites in the vector (Fig 5B, 1^st^ lane, middle panel). In the presence of just P1 nuclease, a similar low level of spontaneous unwinding is observed, resulting in the appearance of two additional faint bands, one between 1650 and 3000 bp and other between 1000 and 1650 bp (Fig. 5B). The addition of DnaA-6xHis results in an increase in intensity of both these bands in a dose dependent manner (Fig. 5A, middle panel, red arrows).

The ScaI digestions of pAP205 show a complex pattern, which we attribute to partially incomplete digestion under the conditions used, and which we have not been able to fully resolve. The most relevant observation is a clearly visible band of between 650 and 850 bp in the presence of both P1 and DnaA-6xHis (Fig. 5A, lower panel, red arrow). We do not observe spontaneous unwinding in the presence of only P1 nuclease, although the pattern is distinct from that of the control lane (Fig 5B, first lane, lower panel).

The DnaA-dependent appearance of the ~2000 bp band in the BglII digest, the ~1200 and ~2200bp bands in the NotI digest, and the ~700 bp band in the ScaI digest localize the DnaA-dependent unwinding of the *C. difficile oriC* in the *oriC2* region (Fig. 5B, gray rectangle, DUE). Moreover, these results suggest that *C. difficile* has a bipartite origin of replication, as successful DnaA-dependent unwinding of *C. difficile* in the *oriC2* region requires both *oriC* regions *(oriC1* and *oriC2).*

### 3.5 Conservation of the origin organisation in related Clostridia

Our results suggest that the origin organization of *C. difficile* resembles that of a more distantly related Firmicute, *B. subtilis.* To extend our observations, we evaluated the genomic organization of the *oriC* region in different organisms phylogenetically related to *C. difficile.* We followed a similar approach as described above for *C. difficile 630Δerm,* taking advantage of the DoriC 10.0 database (http://tubic.tju.edu.cn/doric/public/index.php) (Luo and Gao, 2019). Importantly, our results with respect to the *C. difficile* origin of replication described above were largely congruent with the DoriC 10.0 database (data not shown). We retrieved the predicted *oriC* regions from the DoriC 10.0 database and performed an in-depth analysis of these regions for the closely related *C. difficile* strain R20291 (NC_013316.1), as well as the more distantly related *C. botulinum* A Hall (NC_009698.1), *C. sordelli* AM370 (NZ_CP014150), *C. acetobutylicum* DSM 1731 (NC_015687.1), *C. perfringens* str.13 (NC_003366.1) and *C. tetani* E88 (NC_004557.1) (Table 1).

Similar to *C. difficile 630Δerm,* the genomic context of the origin contains the *rpmH- dnaA-dnaN* region for most of the clostridia selected and mirrors that of *B. subtilis* (Fig. 6). The only exception is *C. tetani* E88 where the uncharacterized CLOTE0041 gene lies upstream of the *dnaA-dnaN* cluster (Fig. 6).

**Figure 6.**
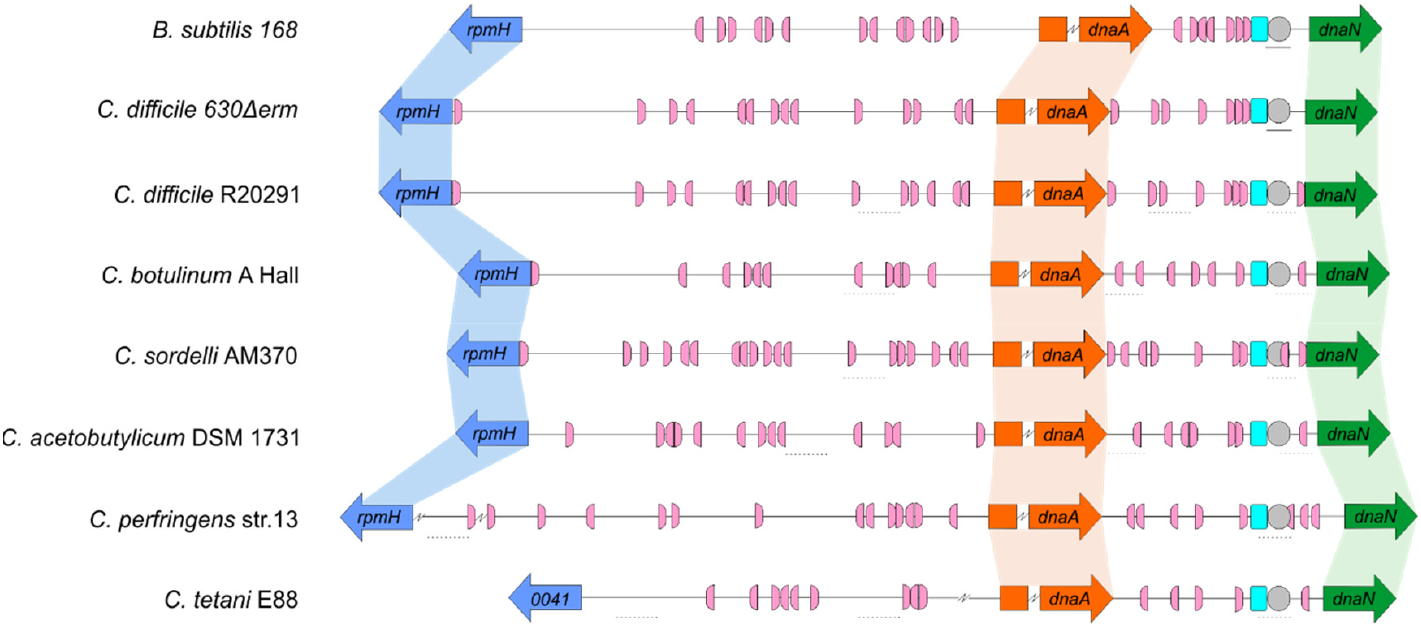
Comparison of the clostridia *oriC* regions. Representation of the origin region and genomic context of B. subtilis, *C. difficile 630Δerm* chromosome and the predicted regions for *C. difficile* R20291, *C. botulinum A* Hall, *C. sordelli* AM370, *C. acetobutylicum* DSM 1731, *C. perfringens* str.13, *C. tetani* E88 (see Table 1). The *rpmH* (blue arrow), *dnaA* (orange arrow) and *dnaN* (green arrow) genes are indicated. Predicted DnaA-boxes are indicated by pink boxes and orientation on the leading (right) and lagging strand (left) are shown. Identification of the experimentally identified unwinding AT-rich regions (lines) and the SIDD-predicted helical instability are shown (dashed lines). The putative DUE is denoted (grey circle). Possible DnaA-trio sequences are shown in light blue boxes. See Material and Methods for detailed information. Alignment of the represented chromosomal regions is based on the location of the DnaA-trio.

We also identified the possible DnaA boxes for the selected clostridia (Fig. 6, pink semicircle). Across the analyzed clostridia, *oriC1* region presented more variability in the number of putative DnaA boxes, from 9 to 19, whereas *oriC2* contained 5 to 9 DnaA boxes, with *C. tetani* E88 with the lowest number of possible DnaA boxes, both at the *oriC1* (9 boxes) and *oriC2* (5 boxes) regions (Fig. 6, pink semi-circle). In all the organisms we observe at least 1 DnaA cluster in each origin region, as also observed for *C. difficile* 630Δ*erm*.

Prediction of DUEs using the SIST program (Zhabinskaya et al., 2015) identified several helically unstable regions that are candidate sites for unwinding (Fig. 6, dashed lines, and Fig. S3). Notably, in all cases one such region in *oriC2* (Fig. 6, grey circle) is preceded immediately by the manually identified DnaA-trio (Fig. 6, light blue circle). Based on our experimental data for *C. difficile 630Δerm,* we suggest that in all analyzed clostridia, DnaA-dependent unwinding occurs at a conserved DUE downstream of the DnaA-trio in the *oriC2* region (Fig. 6).

## 4. Discussion

Chromosomal replication is an essential process for the survival of the cell. In most bacteria DnaA protein is the initiator protein for replication and through a cascade of events leads to the successful loading of the replication complex onto the origin of replication (Mott and Berger, 2007).

Initial characterization of bacterial replication has been assessed in the model organisms *E. coli* and *B. subtilis* (Jameson and Wilkinson, 2017). Despite the similarities the structure of the replication origins and the regulation mechanisms are variable among bacteria (Wolanski et al., 2014). In contrast to *E. coli, B. subtilis* origin region is bipartite, with two intergenic regions upstream and downstream the *dnaA* gene. In *C. difficile* the genomic organization in the predicted cluster *rnpA-rpmH-dnaA-dnaN,* and the presence of AT-rich sequences in the intergenic regions is consistent with a bipartite origin, as in *B. subtilis* (Fig. 3).

The origin region contains several DnaA-boxes with different properties that are recognized by the DnaA protein. The specific binding of DnaA to the DnaA-boxes is mediated mainly through domain IV of the DnaA protein. From DNA bound structures of DnaA it was possible to identify several residues involved in the contact with the DnaA boxes, some of which confer specificity (Blaesing et al., 2000; Fujikawa et al., 2003; Tsodikov and Biswas, 2011). Analysis of the of *C. difficile* DnaA homology in domain IV did not show any difference in the residues involved on the DnaA-box specificity (Fig. 1, vertical arrows), suggesting the same consensus motif conservation as the DnaA-box TTWTNCACA for *E.coli* (Schaper and Messer, 1995). The conserved DnaA-box motif allowed us to identify several DnaA boxes along the intergenic regions of the *oriC.* Like in the bipartite origin of *B. subtilis,* we identified at least one cluster of DnaA-boxes in the *C. difficile oriC,* present at the *oriC1* and the in *oriC2* regions (Fig. 4 and 6). However accurate determination of the *C. difficile* DnaA-boxes was not resolved and further footprinting assays could provide insights on the DnaA-box conservation and affinities. Moreover, it remains to be determined whether the DnaA boxes are crucial for origin firing and/or transcriptional regulation.

The P1 nuclease assays place a region in which DnaA-dependent unwinding occurs in the *oriC2* region of *C. difficile,* supported by the presence of the several features on the *oriC2,* such as the identified DUE and DnaA-trio, both required for unwinding (Kowalski and Eddy, 1989; Richardson et al., 2016). The presence of both *oriC* regions *(oriC1* and *oriC2)* is required for melting *in vitro,* as observed for other bipartite origins (Wolanski et al., 2014). In contrast to the bipartite origin identified in *H. pylori* (Donczew et al., 2012), we did not observe unwinding of the *oriC2* region alone. Though this may be a specific aspect of *C. difficile oriC2,* we cannot exclude that differences in the experimental setup (e.g. DnaA protein purification) could affect these observations. Nevertheless, our data are consistent with DnaA binding the DnaA-box clusters in both *oriC* regions, leading to potential DnaA oligomerization, loop formation, and unwinding at the AT-rich DUE site.

When analyzing the origin region between different clostridia, features similar to those of *C. difficile* are observed, such as conservation of DnaA-box clusters within both *oriC* regions in the vicinity of the *dnaA* gene. Similar to *C. difficile* and *B. subtilis,* a putative DUE element, preceded by the DnaA-trio, was also located within the *oriC2* region (Fig. 4 and 6). Thus, the overall origin organization and mechanism of DNA replication initiation is likely to be conserved within the Firmicutes (Briggs et al., 2012). As spacing of the DnaA-boxes are determinants for the species-specific effective replication (Zawilak et al., 2003; Zawilak-Pawlik et al., 2005), these similarities do no exclude the possibilities that subtle differences in replication initiation exist, and further studies are required.

Additionally, several proteins can interact with the *oriC* region or DnaA, including YabA, Rok, DnaD/DnaB, Soj and HU (Briggs et al., 2012; Jameson and Wilkinson, 2017). In doing so they shape the origin conformation and/or stabilize the DnaA filament or the unwound region, consequently affecting replication initiation.

YabA or Rok affect *B. subtilis* replication initiation (Goranov et al., 2009; Schenk et al., 2017; Seid et al., 2017), but no homologs of these proteins have been identified in *C. difficile.* In *B. subtilis,* DnaD, DnaB and DnaI helicase loader proteins associate sequentially with the origin region resulting in the recruitment of the DnaC helicase protein (Marsin et al., 2001; Velten et al., 2003; Smits et al., 2010; Jameson and Wilkinson, 2017). In *B. subtilis,* DnaD binds to DnaA and it is postulated that this affects the stability of the DnaA filament and consequently the unwinding of the *oriC* (Ishigo- Oka et al., 2001; Martin et al., 2018; Matthews and Simmons, 2019). *B. subtilis* DnaB protein also affects the DNA topology and has been shown to be important for recruiting *oriC* to the membrane (Rokop et al., 2004; Zhang et al., 2005). *C. difficile* lacks a homologue for the DnaB protein, although the closest homolog of the DnaD protein (CD3653) (van Eijk et al., 2017) may perform similar functions in the origin remodeling (van Eijk et al., 2016). Direct interaction of DnaA-DnaD through the DnaA domain I was structurally determined and the residues present at the interface were solved (Martin et al., 2018). Despite high variability of this domain between organisms, half of the identified contacts for the DnaA-DnaD interaction are conserved within *C. difficile,* the S22 (S23 in *B. subtilis* DnaA), T25 (T26), F48 (F49), D51 (D52) and L68 (L69) (Fig. 1) (Martin et al., 2018; Matthews and Simmons, 2019). This might suggest a similar interaction surface for CD3653 on *C. difficile* DnaA. A characterization of the putative interaction between CD3653 and DnaA, and the resulting effect on DnaA oligomerization and origin melting awaits purification and functional characterization of CD3653. The Soj protein, also involved in chromosome segregation, has been shown to interact with DnaA via domain III, regulating DnaA-filament formation (Scholefield et al., 2012) and *C. difficile* encodes at least one uncharacterized Soj homolog. Bacterial histone-like proteins (such as HU and HBsu) can modulate DNA topology and have been shown the influence on *oriC* unwinding and replication initiation in other organisms (Krause et al., 1997; Chodavarapu et al., 2008). *C. difficile* encodes a homologue of HU, HupA (Oliveira Paiva et al., 2019). Though the role of Soj and HupA in DNA replication remains to be elucidated, our experiments show they are not strictly required for origin unwinding. Finally, Spo0A, the master regulator of sporulation, binds to several Spo0A-boxes present in this the *oriC* region in *B. subtilis* (Boonstra et al., 2013). Some of the Spo0A-boxes partially overlap with DnaA-boxes and binding of Spo0A can prevent the DnaA-mediated unwinding, thus playing a significant role on the coordination of between cell replication and sporulation (Boonstra et al., 2013). In *C. difficile,* Spo0A-binding has previously been investigated (Rosenbusch et al., 2012), but a role in DNA replication has not been assessed. For all the regulators with a *C. difficile* homolog discussed above (i.e. CD3653, Soj, HupA and Spo0A), further studies can be envisioned employing the P1 nuclease assays described here to assess the effects on DnaA-mediated unwinding of the origin.

In summary, through a combination of different *in silico* predictions and *in vitro* studies, we have shown DnaA-dependent unwinding in the *dnaA-dnaN* intergenic region of the bipartite *C. difficile* origin of replication. We have analysed the putative origin of replication in different clostridia and observed a conserved organization throughout the Firmicutes, although different mechanisms and modes of regulation might drive the initiation of replication. The present study is the first to characterize the origin region of *C. difficile* and form the start to further unravel the mechanism behind the DnaA-dependent regulation of *C. difficile* initiation of replication.

## Author contributions

AMOP and WKS designed experiments. AMOP and CW performed the *in silico* analyses. AMOP, EVE and AF performed experiments. AMOP and WKS analysed data and wrote the manuscript. All authors read and approved the final version for submission.

## Acknowledgments

We thank Alan Grossman for kindly providing the pAV13 vector and *E. coli* strain CYB1002. We thank Anna Zawilak-Pawlik for kindly providing the pori1ori2 vector and expert help in setting up the P1 assays. We also thank Luís Sousa for help with the SIDD and Pattern Locator coding files.

## References

Abe, Y., Jo, T., Matsuda, Y., Matsunaga, C., Katayama, T., and Ueda, T. (2007). Structure and function of DnaA N-terminal domains: specific sites and mechanisms in inter-DnaA interaction and in DnaB helicase loading on oriC. J Biol Chem 282(24), 17816–17827. doi: 10.1074/jbc.M701841200.

Bawono, P., and Heringa, J. (2014). PRALINE: a versatile multiple sequence alignment toolkit. Methods Mol Biol 1079, 245–262. doi: 10.1007/978-1-62703-646-7_16.

Bazin, A., Cherrier, M.V., Gutsche, I., Timmins, J., and Terradot, L. (2015). Structure and primase-mediated activation of a bacterial dodecameric replicative helicase. Nucleic Acids Res. doi: 10.1093/nar/gkv792.

Blaesing, F., Weigel, C., Welzeck, M., and Messer, W. (2000). Analysis of the DNA-binding domain of Escherichia coli DnaA protein. Mol Microbiol 36(3), 557–569. doi: 10.1046/j.1365-2958.2000.01881.x.

Bleichert, F., Botchan, M.R., and Berger, J.M. (2017). Mechanisms for initiating cellular DNA replication. Science 355(6327). doi: 10.1126/science.aah6317.

Boonstra, M., de Jong, I.G., Scholefield, G., Murray, H., Kuipers, O.P., and Veening, J.W. (2013). Spo0A regulates chromosome copy number during sporulation by directly binding to the origin of replication in Bacillus subtilis. Mol Microbiol 87(4), 925–938. doi: 10.1111/mmi.12141.

Briggs, G.S., Smits, W.K., and Soultanas, P. (2012). Chromosomal replication initiation machinery of low-G+C-content Firmicutes. J Bacteriol 194(19), 5162–5170. doi: 10.1128/JB.00865-12.

Cho, E., Ogasawara, N., and Ishikawa, S. (2008). The functional analysis of YabA, which interacts with DnaA and regulates initiation of chromosome replication in Bacillus subtils. Genes Genet Syst 83(2), 111–125. doi: 10.1266/ggs.83.111.

Chodavarapu, S., Felczak, M.M., Yaniv, J.R., and Kaguni, J.M. (2008). Escherichia coli DnaA interacts with HU in initiation at the E. coli replication origin. Mol Microbiol 67(4), 781–792. doi: 10.1111/j.1365-2958.2007.06094.x.

Chodavarapu, S., and Kaguni, J.M. (2016). Replication Initiation in Bacteria. Enzymes 39, 1–30. doi: 10.1016/bs.enz.2016.03.001.

Crobach, M.J.T., Vernon, J.J., Loo, V.G., Kong, L.Y., Pechine, S., Wilcox, M.H., et al. (2018). Understanding Clostridium difficile Colonization. Clin Microbiol Rev 31(2). doi: 10.1128/CMR.00021-17.

Davey, M.J., and O’Donnell, M. (2003). Replicative helicase loaders: ring breakers and ring makers. Current Biology 13(15), R594–R596. doi: 10.1016/s0960-9822(03)00523-2.

Donczew, R., Weigel, C., Lurz, R., Zakrzewska-Czerwinska, J., and Zawilak-Pawlik, A. (2012). Helicobacter pylori oriC--the first bipartite origin of chromosome replication in Gram-negative bacteria. Nucleic Acids Res 40(19), 9647–9660. doi: 10.1093/nar/gks742.

Ekundayo, B., and Bleichert, F. (2019). Origins of DNA replication. PLoS Genet 15(9), e1008320. doi: 10.1371/journal.pgen.1008320.

Erzberger, J.P., Mott, M.L., and Berger, J.M. (2006). Structural basis for ATP-dependent DnaA assembly and replication-origin remodeling. Nat Struct Mol Biol 13(8), 676–683. doi: 10.1038/nsmb1115.

Erzberger, J.P., Pirruccello, M.M., and Berger, J.M. (2002). The structure of bacterial DnaA: implications for general mechanisms underlying DNA replication initiation. EMBO J 21(18), 4763–4773.

Fossum, S., De Pascale, G., Weigel, C., Messer, W., Donadio, S., and Skarstad, K. (2008). A robust screen for novel antibiotics: specific knockout of the initiator of bacterial DNA replication. FEMS Microbiol Lett 281(2), 210–214. doi: 10.1111/j.1574-6968.2008.01103.x.

Fujikawa, N., Kurumizaka, H., Nureki, O., Terada, T., Shirouzu, M., Katayama, T., et al. (2003). Structural basis of replication origin recognition by the DnaA protein. Nucleic Acids Res 31(8), 2077–2086.

Goranov, A.I., Breier, A.M., Merrikh, H., and Grossman, A.D. (2009). YabA of Bacillus subtilis controls DnaA-mediated replication initiation but not the transcriptional response to replication stress. Mol Microbiol 74(2), 454–466. doi: 10.1111/j.1365-2958.2009.06876.x.

Grimwade, J.E., and Leonard, A.C. (2017). Targeting the Bacterial Orisome in the Search for New Antibiotics. Front Microbiol 8, 2352. doi: 10.3389/fmicb.2017.02352.

Ishigo-Oka, D., Ogasawara, N., and Moriya, S. (2001). DnaD protein of Bacillus subtilis interacts with DnaA, the initiator protein of replication. J Bacteriol 183(6), 2148–2150. doi: 10.1128/JB.183.6.2148-2150.2001.

Jameson, K.H., Rostami, N., Fogg, M.J., Turkenburg, J.P., Grahl, A., Murray, H., et al. (2014). Structure and interactions of the Bacillus subtilis sporulation inhibitor of DNA replication, SirA, with domain I of DnaA. Mol Microbiol 93(5), 975–991. doi: 10.1111/mmi.12713.

Jameson, K.H., and Wilkinson, A.J. (2017). Control of Initiation of DNA Replication in Bacillus subtilis and Escherichia coli. Genes (Basel) 8(1). doi: 10.3390/genes8010022.

Jaworski, P., Donczew, R., Mielke, T., Thiel, M., Oldziej, S., Weigel, C., et al. (2016). Unique and Universal Features of Epsilonproteobacterial Origins of Chromosome Replication and DnaA-DnaA Box Interactions. Front Microbiol 7, 1555. doi: 10.3389/fmicb.2016.01555.

Katayama, T., Kasho, K., and Kawakami, H. (2017). The DnaA Cycle in Escherichia coli: Activation, Function and Inactivation of the Initiator Protein. Front Microbiol 8, 2496. doi: 10.3389/fmicb.2017.02496.

Katayama, T., Ozaki, S., Keyamura, K., and Fujimitsu, K. (2010). Regulation of the replication cycle: conserved and diverse regulatory systems for DnaA and oriC. Nat Rev Microbiol 8(3), 163–170. doi: 10.1038/nrmicro2314.

Kawakami, H., Keyamura, K., and Katayama, T. (2005). Formation of an ATP-DnaA-specific initiation complex requires DnaA Arginine 285, a conserved motif in the AAA+ protein family. J Biol Chem 280(29), 27420–27430. doi: 10.1074/jbc.M502764200.

Kelley, L.A., Mezulis, S., Yates, C.M., Wass, M.N., and Sternberg, M.J. (2015). The Phyre2 web portal for protein modeling, prediction and analysis. Nat Protoc 10(6), 845–858. doi: 10.1038/nprot.2015.053.

Kim, J.S., Nanfara, M.T., Chodavarapu, S., Jin, K.S., Babu, V.M.P., Ghazy, M.A., et al. (2017). Dynamic assembly of Hda and the sliding clamp in the regulation of replication licensing. Nucleic Acids Res 45(7), 3888–3905. doi: 10.1093/nar/gkx081.

Kowalski, D., and Eddy, M.J. (1989). The DNA unwinding element: a novel, cis-acting component that facilitates opening of the Escherichia coli replication origin. EMBO J 8(13), 4335–4344.

Krause, M., Ruckert, B., Lurz, R., and Messer, W. (1997). Complexes at the replication origin of Bacillus subtilis with homologous and heterologous DnaA protein. J Mol Biol 274(3), 365–380. doi: 10.1006/jmbi.1997.1404.

Lawson, P.A., Citron, D.M., Tyrrell, K.L., and Finegold, S.M. (2016). Reclassification of Clostridium difficile as Clostridioides difficile (Hall and O’Toole 1935) Prevot 1938. Anaerobe 40, 95–99. doi: 10.1016/j.anaerobe.2016.06.008.

Luo, H., and Gao, F. (2019). DoriC 10.0: an updated database of replication origins in prokaryotic genomes including chromosomes and plasmids. Nucleic Acids Res 47(D1), D74–D77. doi: 10.1093/nar/gky1014.

Mackiewicz, P., Zakrzewska-Czerwinska, J., Zawilak, A., Dudek, M.R., and Cebrat, S. (2004). Where does bacterial replication start? Rules for predicting the oriC region. Nucleic Acids Res 32(13), 3781–3791. doi: 10.1093/nar/gkh699.

Majka, J., Messer, W., Schrempf, H., and Zakrzewska-Czerwinska, J. (1997). Purification and characterization of the Streptomyces lividans initiator protein DnaA.

Marsin, S., McGovern, S., Ehrlich, S.D., Bruand, C., and Polard, P. (2001). Early steps of Bacillus subtilis primosome assembly. J Biol Chem 276(49), 45818–45825. doi: 10.1074/jbc.M101996200.

Martin, E., Williams, H.E.L., Pitoulias, M., Stevens, D., Winterhalter, C., Craggs, T.D., et al. (2018). DNA replication initiation in Bacillus subtilis: structural and functional characterization of the essential DnaA-DnaD interaction. Nucleic Acids Res. doi: 10.1093/nar/gky1220.

Matthews, L.A., and Simmons, L.A. (2019). Cryptic protein interactions regulate DNA replication initiation. Mol Microbiol 111(1), 118–130. doi: 10.1111/mmi.14142.

Moriya, S., Fukuoka, T., Ogasawara, N., and Yoshikawa, H. (1988). Regulation of initiation of the chromosomal replication by DnaA-boxes in the origin region of the Bacillus subtilis chromosome. EMBO J 7(9), 2911–2917.

Mott, M.L., and Berger, J.M. (2007). DNA replication initiation: mechanisms and regulation in bacteria. Nat Rev Microbiol 5(5), 343–354. doi: 10.1038/nrmicro1640.

Mrazek, J., and Xie, S. (2006). Pattern locator: a new tool for finding local sequence patterns in genomic DNA sequences. Bioinformatics 22(24), 3099–3100. doi: 10.1093/bioinformatics/btl551.

Murray, H., and Koh, A. (2014). Multiple regulatory systems coordinate DNA replication with cell growth in Bacillus subtilis. PLoS Genet 10(10), e1004731. doi: 10.1371/journal.pgen.1004731.

Natrajan, G., Noirot-Gros, M.F., Zawilak-Pawlik, A., Kapp, U., and Terradot, L. (2009). The structure of a DnaA/HobA complex from Helicobacter pylori provides insight into regulation of DNA replication in bacteria. Proc Natl Acad Sci U S A 106(50), 21115–21120. doi: 10.1073/pnas.0908966106.

Necsulea, A., and Lobry, J.R. (2007). A new method for assessing the effect of replication on DNA base composition asymmetry. Mol Biol Evol 24(10), 2169–2179. doi: 10.1093/molbev/msm148.

Nowaczyk-Cieszewska, M., Zyla-Uklejewicz, D., Noszka, M., Jaworski, P., Mielke, T., and Zawilak-Pawlik, A.M. (2019). The role of Helicobacter pylori DnaA domain I in orisome assembly on a bipartite origin of chromosome replication. Mol Microbiol. doi: 10.1111/mmi.14423.

Nozaki, S., and Ogawa, T. (2008). Determination of the minimum domain II size of Escherichia coli DnaA protein essential for cell viability. Microbiology 154(Pt 11), 3379–3384. doi: 10.1099/mic.0.2008/019745-0.

O’Donnell, M., Langston, L., and Stillman, B. (2013). Principles and concepts of DNA replication in bacteria, archaea, and eukarya. Cold Spring Harb Perspect Biol 5(7). doi: 10.1101/cshperspect.a010108.

Ogasawara, N., Moriya, S., von Meyenburg, K., Hansen, F.G., and Yoshikawa, H. (1985). Conservation of genes and their organization in the chromosomal replication origin region of Bacillus subtilis and Escherichia coli. EMBO J 4(12), 3345–3350.

Ogasawara, N., and Yoshikawa, H. (1992). Genes and their organization in the replication origin region of the bacterial chromosome. Mol Microbiol 6(5), 629–634. doi: 10.1111/j.1365-2958.1992.tb01510.x.

Oliveira Paiva, A.M., Friggen, A.H., Qin, L., Douwes, R., Dame, R.T., and Smits, W.K. (2019). The Bacterial Chromatin Protein HupA Can Remodel DNA and Associates with the Nucleoid in Clostridium difficile. J Mol Biol 431(4), 653–672. doi: 10.1016/j.jmb.2019.01.001.

Ozaki, S., and Katayama, T. (2012). Highly organized DnaA-oriC complexes recruit the single-stranded DNA for replication initiation. Nucleic Acids Res 40(4), 1648–1665. doi: 10.1093/nar/gkr832.

Ozaki, S., Kawakami, H., Nakamura, K., Fujikawa, N., Kagawa, W., Park, S.Y., et al. (2008). A common mechanism for the ATP-DnaA-dependent formation of open complexes at the replication origin. J Biol Chem 283(13), 8351–8362. doi: 10.1074/jbc.M708684200.

Ozaki, S., Noguchi, Y., Hayashi, Y., Miyazaki, E., and Katayama, T. (2012). Differentiation of the DnaA-oriC subcomplex for DNA unwinding in a replication initiation complex. J Biol Chem 287(44), 37458–37471. doi: 10.1074/jbc.M112.372052.

Patel, M.J., Bhatia, L., Yilmaz, G., Biswas-Fiss, E.E., and Biswas, S.B. (2017). Multiple conformational states of DnaA protein regulate its interaction with DnaA boxes in the initiation of DNA replication. Biochim Biophys Acta. doi: 10.1016/j.bbagen.2017.06.013.

Richardson, T.T., Harran, O., and Murray, H. (2016). The bacterial DnaA-trio replication origin element specifies single-stranded DNA initiator binding. Nature 534(7607), 412–416. doi: 10.1038/nature17962.

Richardson, T.T., Stevens, D., Pelliciari, S., Harran, O., Sperlea, T., and Murray, H. (2019). Identification of a basal system for unwinding a bacterial chromosome origin. EMBO J 38(15), e101649. doi: 10.15252/embj.2019101649.

Rokop, M.E., Auchtung, J.M., and Grossman, A.D. (2004). Control of DNA replication initiation by recruitment of an essential initiation protein to the membrane of Bacillus subtilis. Mol Microbiol 52(6), 1757–1767. doi: 10.1111/j.1365-2958.2004.04091.x.

Rosenbusch, K.E., Bakker, D., Kuijper, E.J., and Smits, W.K. (2012). C. difficile 630Derm Spo0A Regulates Sporulation, but Does Not Contribute to Toxin Production, by Direct High-Affinity Binding to Target DNA. PloS One. doi: 10.1371/journal.pone.0048608.

Sambrook, J., Fritsch, E.F., and Maniatis, T. (1989). Molecular cloning: a laboratory manual. Cold Spring Harbor, N.Y.: Cold Spring Harbor Laboratory.

Saxena, R., Fingland, N., Patil, D., Sharma, A.K., and Crooke, E. (2013). Crosstalk between DnaA protein, the initiator of Escherichia coli chromosomal replication, and acidic phospholipids present in bacterial membranes. Int J Mol Sci 14(4), 8517–8537. doi: 10.3390/ijms14048517.

Schaper, S., and Messer, W. (1995). Interaction of the initiator protein DnaA of Escherichia coli with its DNA target. J Biol Chem 270(29), 17622–17626. doi: 10.1074/jbc.270.29.17622.

Schenk, K., Hervas, A.B., Rosch, T.C., Eisemann, M., Schmitt, B.A., Dahlke, S., et al. (2017). Rapid turnover of DnaA at replication origin regions contributes to initiation control of DNA replication. PLoS Genet 13(2), e1006561. doi: 10.1371/journal.pgen.1006561.

Scholefield, G., Errington, J., and Murray, H. (2012). Soj/ParA stalls DNA replication by inhibiting helix formation of the initiator protein DnaA. EMBO J 31(6), 1542–1555. doi: 10.1038/emboj.2012.6.

Scholefield, G., and Murray, H. (2013). YabA and DnaD inhibit helix assembly of the DNA replication initiation protein DnaA. Mol Microbiol 90(1), 147–159. doi: 10.1111/mmi.12353.

Seid, C.A., Smith, J.L., and Grossman, A.D. (2017). Genetic and biochemical interactions between the bacterial replication initiator DnaA and the nucleoid-associated protein Rok in Bacillus subtilis. Mol Microbiol 103(5), 798–817. doi: 10.1111/mmi.13590.

Sekimizu, K., Bramhill, D., and Kornberg, A. (1988). Sequential early stages in the in vitro initiation of replication at the origin of the Escherichia coli chromosome. J Biol Chem 263(15), 7124–7130.

Smits, W.K., Goranov, A.I., and Grossman, A.D. (2010). Ordered association of helicase loader proteins with the Bacillus subtilis origin of replication in vivo. Mol Microbiol 75(2), 452–461. doi: 10.1111/j.1365-2958.2009.06999.x.

Smits, W.K., Lyras, D., Lacy, D.B., Wilcox, M.H., and Kuijper, E.J. (2016). Clostridium difficile infection. Nature Reviews Disease Primers 2, 16020. doi: 10.1038/nrdp.2016.20.

Smits, W.K., Merrikh, H., Bonilla, C.Y., and Grossman, A.D. (2011). Primosomal proteins DnaD and DnaB are recruited to chromosomal regions bound by DnaA in Bacillus subtilis. J Bacteriol 193(3), 640–648. doi: 10.1128/JB.01253-10.

Speck, C., Weigel, C., and Messer, W. (1999). ATP- and ADP-dnaA protein, a molecular switch in gene regulation. EMBO J 18(21), 6169–6176. doi: 10.1093/emboj/18.21.6169.

Sutton, M.D., and Kaguni, J.M. (1997). Novel alleles of the Escherichia coli dnaA gene. J Mol Biol 271(5), 693–703. doi: 10.1006/jmbi.1997.1209.

Torti, A., Lossani, A., Savi, L., Focher, F., Wright, G.E., Brown, N.C., et al. (2011). Clostridium difficile DNA polymerase IIIC: basis for activity of antibacterial compounds. Current Enzyme Inhibition 7(October).

Tsodikov, O.V., and Biswas, T. (2011). Structural and thermodynamic signatures of DNA recognition by Mycobacterium tuberculosis DnaA. J Mol Biol 410(3), 461–476. doi: 10.1016/j.jmb.2011.05.007.

van Eijk, E., Anvar, S.Y., Browne, H.P., Leung, W.Y., Frank, J., Schmitz, A.M., et al. (2015). Complete genome sequence of the Clostridium difficile laboratory strain 630Deltaerm reveals differences from strain 630, including translocation of the mobile element CTn5. BMC Genomics 16, 31. doi: 10.1186/s12864-015-1252-7.

van Eijk, E., Boekhoud, I.M., Kuijper, E.J., Bos-Sanders, I., Wright, G., and Smits, W.K. (2019). Genome Location Dictates the Transcriptional Response to PolC Inhibition in Clostridium difficile. Antimicrob Agents Chemother 63(2). doi: 10.1128/AAC.01363-18.

van Eijk, E., Paschalis, V., Green, M., Friggen, A.H., Larson, M.A., Spriggs, K., et al. (2016). Primase is required for helicase activity and helicase alters the specificity of primase in the enteropathogen Clostridium difficile. Open Biol 6(12). doi: 10.1098/rsob.160272.

van Eijk, E., Wittekoek, B., Kuijper, E.J., and Smits, W.K. (2017). DNA replication proteins as potential targets for antimicrobials in drug-resistant bacterial pathogens. J Antimicrob Chemother 72(5), 1275–1284. doi: 10.1093/jac/dkw548.

Vellanoweth, R.L., and Rabinowitz, J.C. (1992). The influence of ribosome-binding-site elements on translational efficiency in Bacillus subtilis and Escherichia coli in vivo. Mol Microbiol 6(9), 1105–1114. doi: 10.1111/j.1365-2958.1992.tb01548.x.

Velten, M., McGovern, S., Marsin, S., Ehrlich, S.D., Noirot, P., and Polard, P. (2003). A two-protein strategy for the functional loading of a cellular replicative DNA helicase. Mol Cell 11(4), 1009–1020. doi: 10.1016/s1097-2765(03)00130-8.

Warriner, K., Xu, C., Habash, M., Sultan, S., and Weese, S.J. (2017). Dissemination of Clostridium difficile in food and the environment: Significant sources of C. difficile community-acquired infection? J Appl Microbiol 122(3), 542–553. doi: 10.1111/jam.13338.

Weigel, C., Schmidt, A., Seitz, H., Tungler, D., Welzeck, M., and Messer, W. (1999). The N-terminus promotes oligomerization of the Escherichia coli initiator protein DnaA. Mol Microbiol 34(1), 53–66. doi: 10.1046/j.1365-2958.1999.01568.x.

Wolanski, M., Donczew, R., Zawilak-Pawlik, A., and Zakrzewska-Czerwinska, J. (2014). oriC-encoded instructions for the initiation of bacterial chromosome replication. Front Microbiol 5, 735. doi: 10.3389/fmicb.2014.00735.

Xu, W.C., Silverman, M.H., Yu, X.Y., Wright, G., and Brown, N. (2019). Discovery and development of DNA polymerase IIIC inhibitors to treat Gram-positive infections. Bioorg Med Chem 27(15), 3209–3217. doi: 10.1016/j.bmc.2019.06.017.

Zawilak-Pawlik, A., Kois, A., Majka, J., Jakimowicz, D., Smulczyk-Krawczyszyn, A., Messer, W., et al. (2005). Architecture of bacterial replication initiation complexes: orisomes from four unrelated bacteria. Biochem J 389(Pt 2), 471–481. doi: 10.1042/BJ20050143.

Zawilak-Pawlik, A., Nowaczyk, M., and Zakrzewska-Czerwinska, J. (2017). The Role of the N-Terminal Domains of Bacterial Initiator DnaA in the Assembly and Regulation of the Bacterial Replication Initiation Complex. Genes (Basel) 8(5). doi: 10.3390/genes8050136.

Zawilak, A., Durrant, M.C., Jakimowicz, P., Backert, S., and Zakrzewska-Czerwinska, J. (2003). DNA binding specificity of the replication initiator protein, DnaA from Helicobacter pylori. J Mol Biol 334(5), 933–947.

Zhabinskaya, D., Madden, S., and Benham, C.J. (2015). SIST: stress-induced structural transitions in superhelical DNA. Bioinformatics 31(3), 421–422. doi: 10.1093/bioinformatics/btu657.

Zhang, W., Carneiro, M.J., Turner, I.J., Allen, S., Roberts, C.J., and Soultanas, P. (2005). The Bacillus subtilis DnaD and DnaB proteins exhibit different DNA remodelling activities. J Mol Biol 351(1), 66–75. doi: 10.1016/j.jmb.2005.05.065.

Zorman, S., Seitz, H., Sclavi, B., and Strick, T.R. (2012). Topological characterization of the DnaA-oriC complex using single-molecule nanomanipuation. Nucleic Acids Res 40(15), 7375–7383. doi: 10.1093/nar/gks371.

