## Supplemental Material and Figures for "Identification of the unwinding region in the *Clostridioides difficile* chromosomal origin of replication"

### 1. Supplementary Data

**Pattern search:**
NTATCCACA
TNATCCACA
TTANCCACA
TTATCNACA
TTATCCNCA
TTATCCANA
TTATCCACN
TTNTCCACA
TGTGGATAN
TGTGGATNA
TGTGGNTAA
TGTNGATAA
TGNGGATAA
TNTGGATAA
NGTGGATAA
TGTGGANAA
TTWTNCACA
TGTGNAWAA
NTWTNCACA
TGTGNAWAN
TNWTNCACA
TGTGNAWNA
TTWNNCACA
TGTGNNWAA
TTWTNNACA
TGTNNAWAA
TTWTNCNCA
TGNGNAWAA
TTWTNCANA
TNTGNAWAA
TTWTNCACN
NGTGNAWAA

### 2. Supplementary Figures

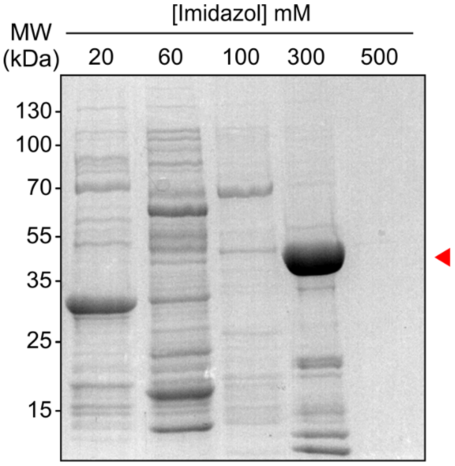


**Figure S1. HisTrap purification of *C. difficile* DnaA-6xHis.** Samples of DnaA-6xHis HisTrap purification from the elution fraction 2 at binding buffer with different imidazole concentrations (20, 60, 100, 300 and 500 mM) were separated by 12% SDS– PAGE and stained with Coomassie brilliant blue. DnaA-6xHis is observed with an approximate molecular weight of 51 kDa (red arrow), and eluted in Binding buffer supplemented with >300 mM imidazole.

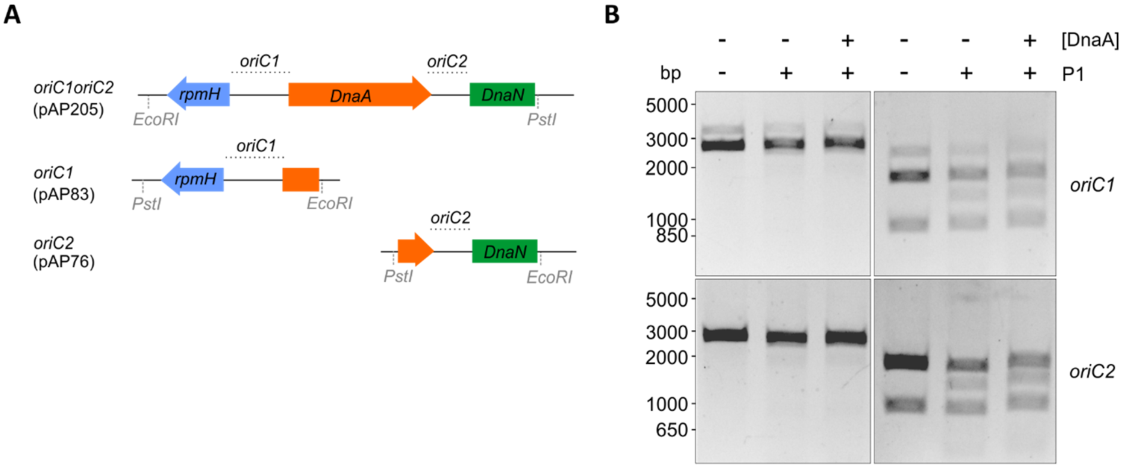


**Figure S2. P1 nuclease assay of the individual *C. difficile oriC regions*. A)** Representation of the *oriC* regions present in the used vectors for P1 nuclease assay, *oriC1oriC2* (pAP205), *oriC1*- (pAP83) and *oriC2* (pAP76)-containing vectors. The predicted *oriC* regions (dotted lines) and included genes are represented, *rpmH* (blue), *dnaA* (orange), and *dnaN* (green). P1 nuclease assay of pAP83 (*oriC1*, upper panel) and pAP76 (*oriC2*, lower panel). Digestion of the vector with the restriction enzymes BglII (left panel) or NotI (right panel). Digestion of the vectors with the restriction enzymes (lanes 1-3). Treatment of the fragments with P1 nuclease only (lane 2) and incubated with 0.14 µM of *C. difficile* DnaA-6xHis protein (lane 3). Higher DnaA-6xHis were tested with same profile (data not shown). The DNA fragments were separated in a 1% agarose gel and analyzed with ethidium bromide staining. Spontaneous unwinding is observed and no DnaA-dependent unwinding is detected.

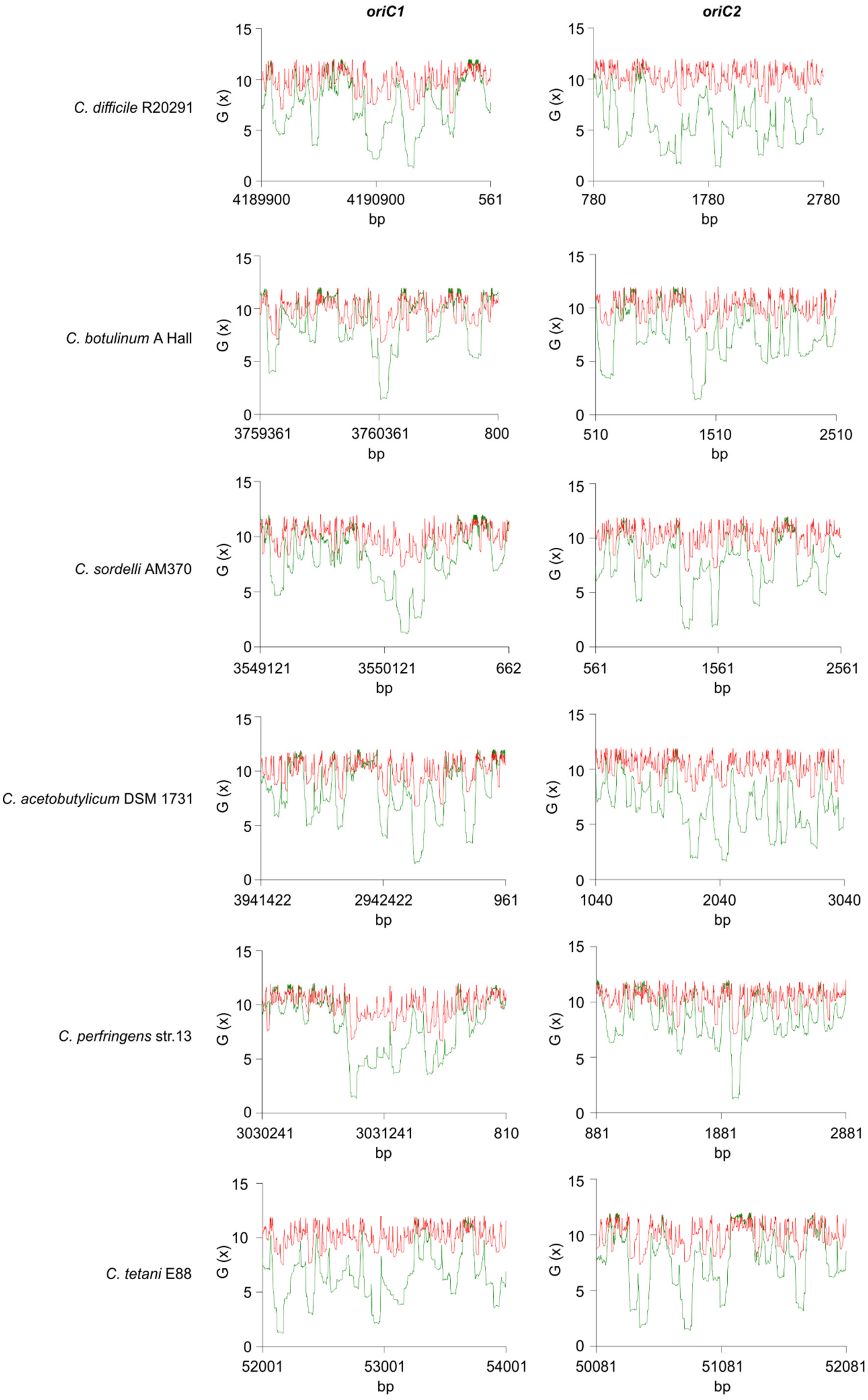
oriC1
oriC2
C. difficile R20291
C. botulinum A Hall
C. sordelli AM370
C. acetobutylicum DSM 1731
C. perfringens str.13
C. tetani E88
G (x)
bp
4189900
4190900
561
780
1780
2780
3759361
3760361
800
510
1510
2510
3549121
3550121
662
561
1561
2561
3941422
2942422
961
1040
2040
3040
3030241
3031241
810
881
1881
2881
52001
53001
54001
50081
51081
52081

**Figure S3. SIDD analysis different clostridia.** Analyis of 2.0 kb fragments comprising *oriC1* and *oriC2* in *C. difficile* R20291, *C. botulinum A* Hall, *C. sordelli* AM370, *C. acetobutylicum* DSM 1731, *C. perfringens* str.13, *C. tetani* E88 (see Table 1 in the main body of the manuscript). Nucleotide positioning is indicated. Predicted free energies G(x) for duplex destabilization at a superhelical density of σ = -0.06 (green) or σ = -0.04 (red).
